# A compact Druantia defense clears phage infections *via* single-stranded DNA recognition and directional duplex unwinding

**DOI:** 10.64898/2026.05.13.724758

**Authors:** Stephanie Himpich, Thomas Gaudin, Lena M. Grass, Huihui Li, Vu Van Loi, Chao Chen, Eberhard Klauck, Philipp F. Popp, Benno Kuropka, Tarek Hilal, Bernhard Loll, Marc Erhardt, Haike Antelmann, Chase L. Beisel, Markus C. Wahl

## Abstract

Bacteria encode diverse anti-phage defense systems triggered by invader-specific molecular cues. Here, we report that the compact type III-A Druantia system recognizes exposed single-stranded DNA to drive phage clearance. Using representative systems from *Escherichia coli*, we show that the two encoded proteins, DruE and DruH, together clear restriction-sensitive or recombination-prone phages without affecting cell growth or viability. DruE dimerizes and engages DNA at exposed single-stranded regions to unwind DNA with 3′-to-5′ directionality, resorting to unique molecular lock, wedge, and clamp elements that aid strand separation and processive translocation. DruH is a monomer in isolation and indirectly interacts with DruE and other host proteins under uninfected conditions, with an infection resulting in the dissociation of the complex. Taken together, our results reveal that exposed single-stranded DNA can trigger bacterial immunity through the directional helicase activity of type III-A Druantia.

**Highlights:**

- The *E. coli* Druantia III-A defense, comprising DruE and DruH, clears infecting phages
- DruE dimers bind exposed single-stranded DNA and unwind the upstream DNA duplex
- 3′-to-5′ DNA unwinding is aided by molecular lock, wedge, and clamp elements
- DruE interacts with DruH and host proteins, which are displaced upon infection

## INTRODUCTION

Bacteria evolved an extensive collection of immune defenses to counteract infections by bacteriophages and other mobile genetic elements^1–3^. Each defensive system is triggered by a molecular cue unique to the invader, ensuring immunity is only activated in infected cells. Numerous cues have been identified, spanning invader-specific components, such as capsid proteins or structured RNAs^4–7^; invader-specific genetic features, such as DNA or RNA sequences or distinct chemical modifications; defined changes in cellular metabolism, such as the depletion of dNTPs ^8,9^; inhibited cellular processes, such as transcription^10^; and inhibited immune defenses, such as through RecBCD^11^. Recognition of these cues then leads the defense to clear the invader or enact growth arrest to prevent dissemination of the invader from the infected cell^12^. Despite the many identified activating cues, other features unique to an active infection that trigger bacterial immunity likely await discovery.

One growing set of known activating cues involves DNA topology unique to an invader. Chemical modifications present or absent in the invader DNA are one common trigger, such as unmethylated DNA cleaved by types I, II, and III restriction-modifications (RM) systems, or DNA with α-glucosyl-5-hydroxymethylcytosines cleaved by the type IV restriction endonuclease GmrSD^13,14^. Separately, free DNA ends absent in circular chromosomal DNA offer another trigger, such as free DNA ends recognized by Shedu, free ends or hairpins recognized by Lamassu, or a 5′ single-stranded (ss) overhang in unmethylated DNA recognized by DISARM, with recognition leading to cleavage of the DNA or, in the case of Lamassu, cellular DNA that drives abortive infection^15–20^. Even when circular, short DNA triggers Wadjet to extrude and then cleave the DNA^21,22^. Finally, a DNA end entering through the cell membrane triggers the membrane-associated defense SNIPE, which degrades the incoming DNA^23^. What remains unclear is whether other aspects of DNA topology can trigger defenses as part of an immune response.

One potential source of new modes of DNA recognition could come from a previously uncharacterized set of defense systems called Druantia^24^. Druantia systems, named after the Celtic goddess and protector of trees^24^, are unified by the shared *druE* gene, encoding a YprA/MrfA-like helicase^25,26^. DruE contains a tandem RecA domain core and has been assigned to helicase superfamily 2 (SF2). In addition to other known helicase auxiliary domains, DruE harbors a MrfA Zn^2+^-binding (MZB), also known as domain of unknown function (DUF) 1998 or Helicase Allosteric Relay (HAR); the latter has been characterized as a zinc-binding domain required for helicase activity in the DNA helicase, mitomycin repair factor A (MrfA)^27^. Homologs of DruE are found in other validated or predicted defenses^25,26^, with the SF2 helicase core and MZB domains separately present in DrmA and DrmB of DISARM^20,28^. The specific composition of DruE and the identity of other encoded proteins divides Druantia systems into three types (I, II, III) and two subtypes (III-A, III-B). Of these systems, III-A and III-B are the most compact, encoding only one additional protein, DruH, of unknown function that lacks annotated domains^24–26^. Representative III-A systems conferred defense in *Escherichia coli* and *Pseudomonas aeruginosa* against a broad range of phages^25,26,29,30^, including T4 phage when lacking glucosyl-5-hydroxymethylcytosines^31^. While the dependency on DNA modifications and the shared homology with the DNA end-triggered defense DISARM suggest that some feature of invader DNA triggers Druantia^20,28^, the molecular details remain to be elucidated.

Here, we report that the compact type III-A Druantia system utilizes ssDNA as a trigger to clear sensitive phages from an infected cell. The system confers defense against diverse phages unified by known sensitivity to DNA restriction or relying on recombinase-based DNA replication. The DruE helicase forms a dimer that binds exposed ssDNA, leading to 3′-to-5′ unwinding of flanking double-stranded regions through a series of defined conformational changes, while DruH is monomeric with a yet-to-be elucidated function. DruE interacts with DruH indirectly under non-infecting conditions, presumably through co-interacting host proteins that contribute to Druantia-mediated defense, while a phage infection causes DruH to disassociate from DruE. In total, these findings establish the compact Druantia III-A system as a DNA-centric defense that clears phages through the recognition of exposed ssDNA and directional unwinding, expanding the known triggers of phage defense and laying a foundation on which to characterize other Druantia types as well as newly discovered DruE-like systems.

## RESULTS

### Druantia III-A clears sensitive phages

To characterize the molecular properties of Druantia type III-A systems, we began with a representative system in *E. coli* ATCC8739 (**Figure 1A**) that, when deleted, sensitizes the strain to multiple phages^25,26^. As the flanking Zorya II and ARMADA II systems exhibit synergy with the Druantia system that could confound elucidating Druantia’s individual contributions ^25,26,32^, we cloned the system’s *druHE* operon under control of its native putative promoter on a low-copy plasmid in the *E. coli* laboratory strain MG1655 that is highly susceptible to infection by diverse phages. We then assessed sensitivity of the strain to the Basel phage collection comprising 68 double-stranded DNA (dsDNA) phages^33^ as well as additional dsDNA phages (**Figure 1B**). Compared to MG1655 strain encoding an empty vector control, the Druantia system conferred resistance against 38 phages from the families *Drexlerviridae, Queuovirinae, Demerecviridae* (specifically the *Eseptimavirus* genus)*, Vequintavirinae*, and *Stephanstirmvirinae* (**Figure 1B**). Phages from the *Tequatrovirus* family were insensitive to the Druantia system, in line with prior work^31^. In contrast to the phage families insensitive to the Druantia system, the sensitive families are also targets of RM-systems, possess a long tail, are prone to recombination through their infection cycle, or combinations thereof^33–35^.

**Figure 1.**
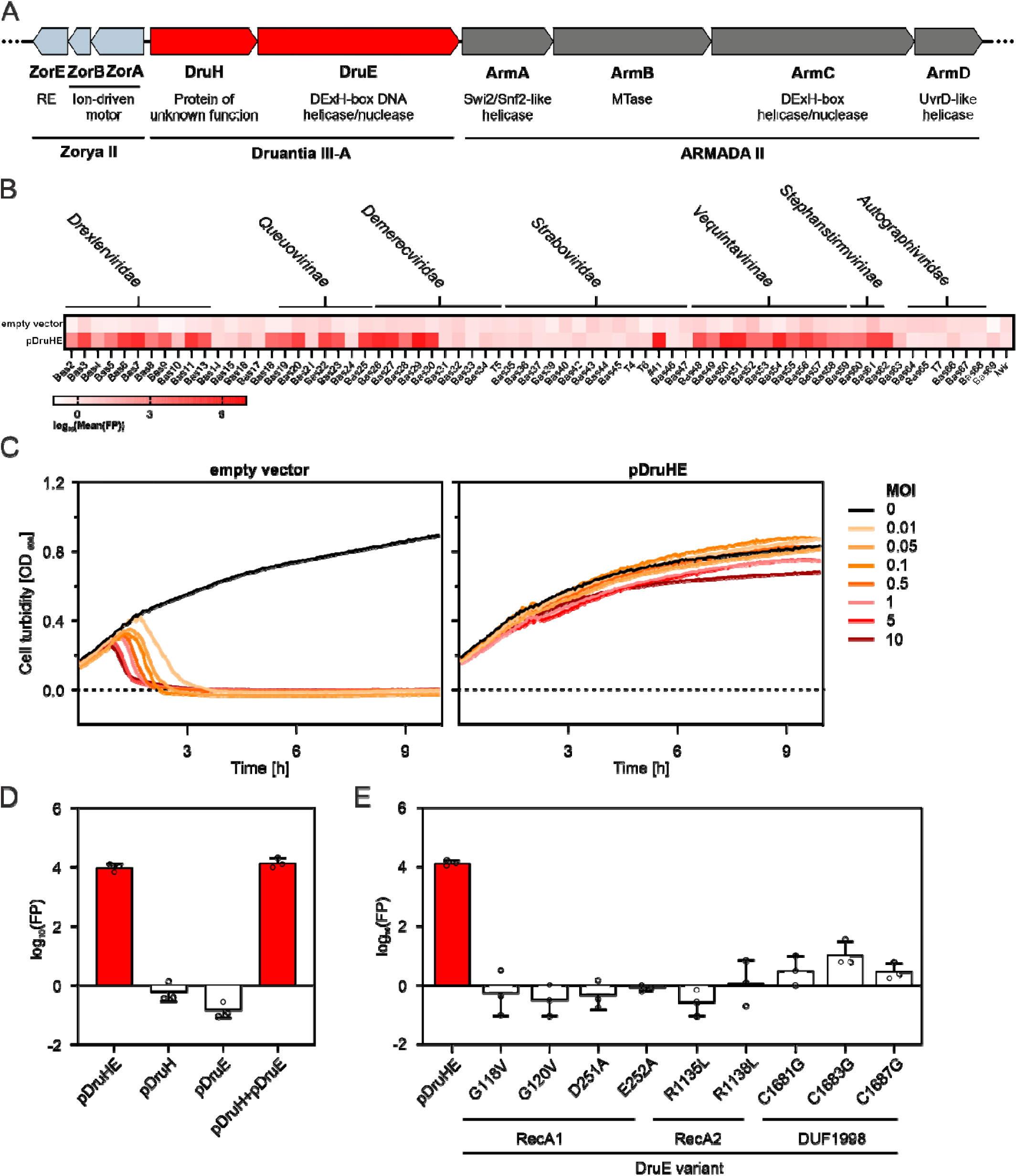
Druantia III-A clears phages that are sensitive to DNA restriction and prone to recombination. (A) Genomic organization of the Druantia III-A system in *E. coli* ATCC8739 and EDL933 with predicted encoded proteins. The system is flanked by two other defenses, a Zorya type II system and an ARMADA type II system, that comprise a satellite phage integrated defensive and ecotypic replicon. (B) Protection conferred by Druantia III-A across the Basel phage collection. The reported fold protection (FP) is based on plaque formation and compares *E. coli* MG1655 with pDruHE to an empty-vector control. Each box represents the mean of log-transformed values from three biological replicates. (C) Protection conferred by Druantia III-A against phage Bas29 in liquid culture. The OD_600_ of a culture of *E. coli* MG1655 with pDruHE or an empty-vector control were monitored over time following the addition of the phage at different MOI. Curves represent the mean of three biological replicates. (D) Essentiality of *druE* and *druH* for anti-phage defense by Druantia III-A against phage #41. The reported fold protection (FP) is based on plaque formation and compares *E. coli* MG1655 with the indicated plasmid to an empty-vector control. To assess gene essentiality, *druE* or *druH* was deleted from pDruHE to form pDruH or pDruE, respectively. Complementation was achieved by combining both plasmids, which contain separate antibiotic resistance markers. Bars and error bars represent the mean ± SD from three biological replicates. Open circles represent values from individual experiments. Red bars indicate defense. (E) Impact of the DruE mutants on anti-phage resistance by Druantia III-A against phage #41. The reported fold protection (FP) is based on plaque formation and compares *E. coli* MG1655 with the indicated plasmid to an empty-vector control. Bars and error bars represent the mean ± SD from three biological replicates. Open circles represent values from individual experiments. Red bars indicate defense.

Phage defenses normally act by clearing the invader, induce growth arrest, or undergo abortive infection that sacrifices the infected cell. After verifying that phages can be adsorbed by the heterologous expression (**Figure S1A**), we determined which mode the Druantia III-A system employs. To do so, we followed the growth of liquid cultures at different multiplicities of infection (MOI) of Bas29, with Druantia conferring over 10^6^-fold protection against this phage (**Figure 1C**). The cultures grew even at an MOI of 10, in line with Druantia clearing the phage. Host DNA methylation was not a requirement for activation or self-protection, as the Druantia system was not cytotoxic in a DNA methylation-free *E. coli* strain^36^ and similarly conferred resistance to Bas29 in this strain (**Figures S1B and S1C**). Thus, the Druantia III-A system acts as a broad phage clearance system independent of the host’s DNA methylation state.

The isolated system further offered an opportunity to affirm the essentiality of both DruE and DruH as well as conserved functional domains. Here, mutants were paired with *Krishvirus* #41, with the full system conferring over 10^4^-fold protection against this phage (**Figure 1B**). Removing either *druE* or *druH* abolished protection, paralleling the impact of gene deletions made in the native host on phage sensitivity^26^, while complementing the deletion with the same gene heterologously expressed from a separate plasmid restored protection (**Figure 1D**). We further introduced disruptive mutations affecting the helicase core of DruE comprising two RecA-like domains (RecA1, RecA2) and the MZB domain. Altering conserved helicase motifs I (GxGKT, G118V/G120V) or II (DExH, D251A/E252A) of RecA1, motif IV (RAGR, R1135L/R1138L) of RecA2, or cysteine residues within the zinc-binding domain of the MZB domain (C1681G/C1683G/C1687G) abolished protection (**Figure 1E**). DruH lacks any annotated domains. Therefore, the Druantia III-A system confers defense through its two proteins, DruE and DruH, and requires the core helicase and MZB domains of DruE.

### DruE is a 3**′**-to-5**′** directional DNA helicase

We next aimed to characterize the properties of DruE contributing to immune defense. We specifically purified DruE encoded in the enterohemorrhagic *E. coli* (EHEC) outbreak strain EDL933, which shares 99% residue identity with DruE from ATCC8739 and also is co-encoded with Zorya II and ARMADA II^26^. DruE was produced *via* a recombinant baculovirus in insect cells and purified to near-homogeneity (**Figure S2B**). In agreement with expectations that DruE is a DNA helicase^24–26^, the purified protein stably associated with a 15-nucleotide (nt) ssDNA (K_d_ = 125 +/- 8 nM) but did not bind ssRNA of the same sequence in a fluorescence anisotropy (FA) assay (**Figure 2A**). Electrophoretic mobility shift assays (EMSAs) with an alternative ssDNA and a corresponding blunt-ended dsDNA substrate confirmed sequence-independent, ss-specific DNA affinity of DruE (**Figure S3A**). DruE also exhibited intrinsic ATPase activity that increased twofold in the presence of ssDNA but not ssRNA (**Figure 2B**). These results show that DruE binds ssDNA, but not ssRNA, that stimulates ATP hydrolysis.

**Figure 2.**
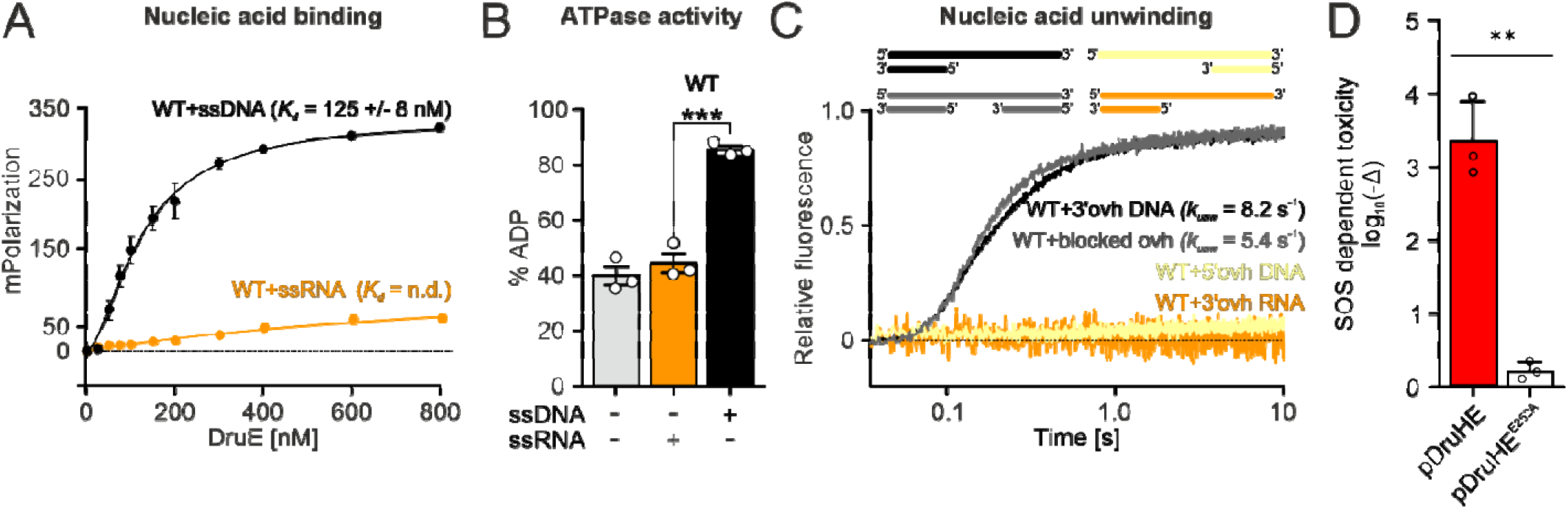
DruE is an efficient DNA helicase that recognizes ssDNA regions. (A) FA assays monitoring binding of DruE to 15-nt ssDNA and 15-nt ssRNA. Data represent means ± SEM of three independent experiments using the same biochemical samples. Here and in following figures, *K_d_* values were derived by fitting the data to a single exponential Hill function (fraction bound = *A*[protein]^n^/([protein]^n^+*K* ^n^); *A*, fitted maximum of nucleic acid bound; n, Hill coefficient; *K_d_*, dissociation constant)^64^. (B) Intrinsic and RNA- or DNA-stimulated ATPase activities of DruE derived from thin-layer chromatographic analyses. Data represent the mean ± SEM of three independent experiments using the same biochemical samples. Significance was assessed *via* unpaired Student’s-t-tests. *, p ≤ 0.05; **, p ≤ 0.01; ns, not significant. (C) Unwinding of 3′ overhang (ovh) DNA, 5′ overhang DNA, blocked overhang DNA, and 3′ overhang RNA by DruE monitored *via* stopped-flow/fluorescence assays. Single representative traces of three independent measurements are shown. Here and in following figures, amplitude-weighted unwinding rate constants were derived as *k_unw_*= ∑(*A_i_k_i_*^2^)/∑(*k_i_A_i_*) by fitting the time traces to a double exponential equation (fraction unwound = *A_fas_*_t_·(1-exp(-*k_fast_t*))+*A_slow_*·(1-exp(-*k_slow_t*)); *A_fast/slow_*, unwinding amplitudes of the fast/slow phases; *k_fast/slow_*, unwinding rate constants of the fast/slow phases [s^-1^]; *t*, time [s]). (D) SOS-dependent toxicity of Druantia III-A in *E. coli* MG1655 exposed to mitomycin C. The relative CFU reduction due to exposure to mitomycin C is reported when comparing the presence of the Druantia III-A system to the empty-vector control. Bars and error bars represent the mean ± SD of the log-transformed values from three biological replicates. Open circles represent individual replicates. Significance was assessed by Welch’s t-tests. **, p < 0.01.

To test if DruE can unwind nucleic acid duplexes, we conducted multi-round, fluorescence-based unwinding assays using a stopped-flow device. DruE efficiently unwound a 12-base pair (bp) DNA duplex containing a 31-nt 3′ overhang, while it failed to unwind the same DNA duplex containing an equal-length 5′ overhang or a corresponding RNA duplex with a 3′ overhang (**Figure 2C**). The free 3′ end was not necessary for DNA unwinding, as DruE efficiently displaced a 12-nt DNA oligo hybridized to the 5′ end of a 48-nt DNA whose 3′ overhang was blocked by another 12-nt complementary oligo (**Figure 2C**). DruE, thus, recognizes ssDNA, including internally within dsDNA, and unwinds flanking dsDNA in the 3′-to-5′ direction.

While DruE recognized ssDNA that could be present in the host chromosome, the immune response of the complete Druantia III-A system was directed to the sensitive phage without impacting host fitness or viability (**Figure 1C**). We hypothesized that inducing ssDNA formation in the host could cause the Druantia system to become cytotoxic through self-recognition. To test this hypothesis, we assessed viability of Druantia-containing cells exposed to the DNA-damaging agent mitomycin C, which induces DNA repair through homologous recombination that forms ssDNA intermediates^37^ (**Figure 2D**). In line with this hypothesis, exposure to mitomycin C reduced colony formation over 10^3^-fold compared to an empty-vector control, whereas introduction of the inactivating DruE E525A substitution abrogated sensitivity to mitomycin C (**Figure 2D**). These results provide further evidence that the Druantia III-A system is triggered by ssDNA absent in cells under normal growth conditions.

### DruE helicase activity relies on unique accessory domains

To elucidate the molecular mechanism of DruE-mediated DNA unwinding, we structurally analyzed DruE in complex with a poorly hydrolyzable ATP analog (ATPүS) and a forked DNA substrate (13-bp duplex region, 17-nt overhangs; **Figures 3A and 3B**). Cryogenic electron microscopy (cryoEM) in combination with single-particle analysis yielded one monomeric and five dimeric constellations, with all DruE protomers bound to ATPγS and DNA (**Figures S5-S7; Table S1**). DruE can be dissected into eleven domains, mostly aligning with those predicted by bioinformatics analyses^25^: two RecA-like domains (RecA1, RecA2), a winged-helix domain (WH), a double Zn-binding domain (dZBD), a duplex-binding domain (DBD), a helical dimerization domain (DD), an extended WH domain (eWH), an OB-fold and Zn-binding domain (OBZ), the MZB domain and two phospholipase D-like domains (PLD1, PLD2; **Figure 3B**). WH, dZBD, DBD and DD are positioned around the flank of the RecA2 domain opposite RecA1. The eWH, OBZ, MZB and PLD1/2 domains are arranged in a circular fashion on top of the RecA1 domain, with the eWH, OBZ and MZB expanding the RecA1/2 core, resembling the DNA helicase MrfA^27^. DruE further contains five zinc-binding motifs (ZBMs; **Figure S8**), with two in the dZBD domain, two in the OBZ domain, and one in the MZB domain. Only one motif in the dZBD and one in the OBZ domain directly contact the bound DNA. The DD could not be resolved in the monomer, as the domain is inserted into the DBD domain *via* two flexible linkers and is presumably dynamically disordered when not bound to a second protomer.

**Figure 3.**
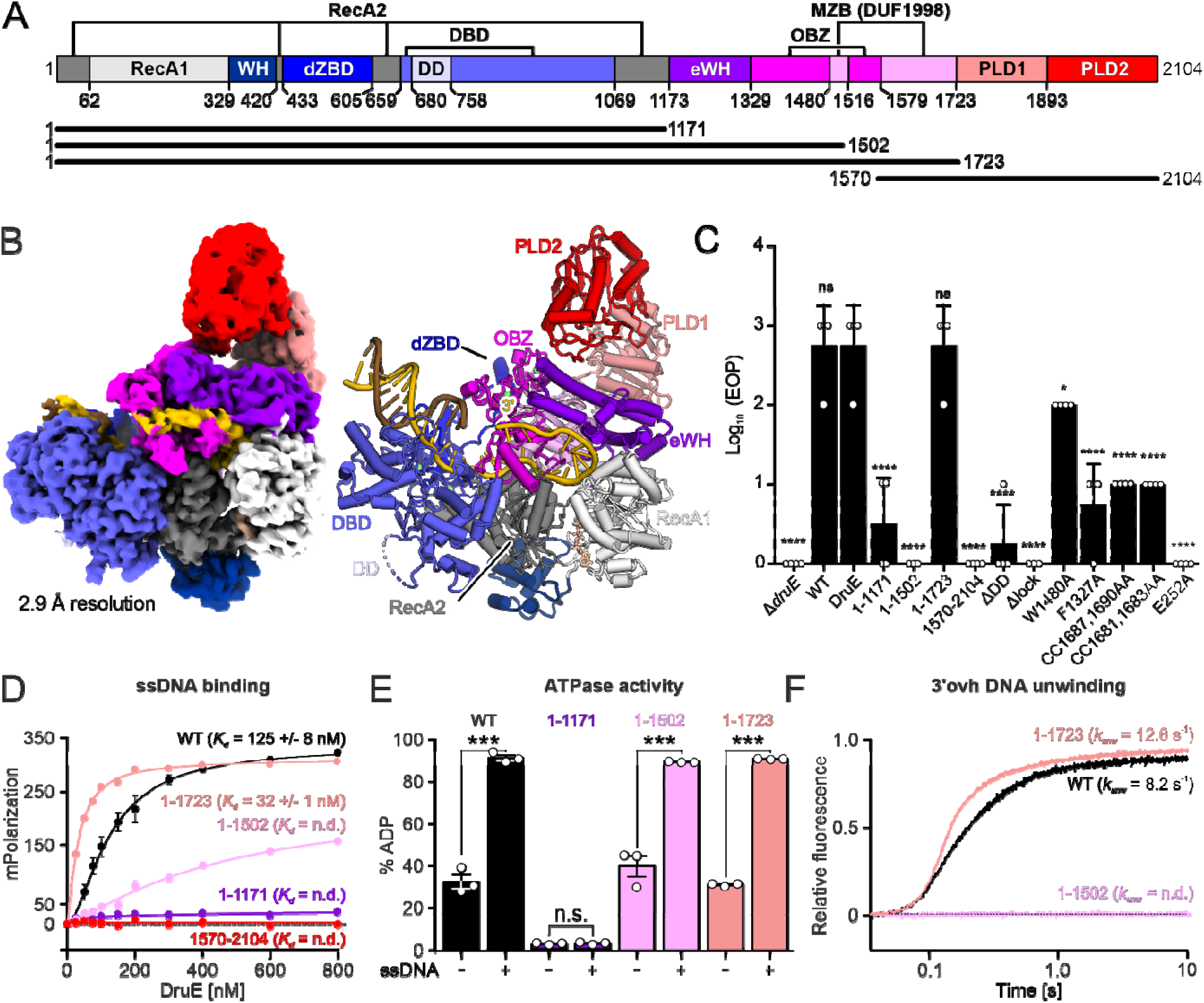
The helicase DruE relies on its MZB domain for duplex unwinding. (A) Domain organization of DruE and schematic representation of DruE truncation variants (DruE^1-1171^, DruE^1-1502^, DruE^1-1723^, and DruE^1570-2104^). (B) CryoEM reconstruction (left) and cartoon representation (right) of monomeric DruE in complex with a forked DNA substrate and the weakly hydrolyzable ATP analog, ATPγS. (C) EOP assays of Bas26 against EDL933 monitoring the complementation of the EDL933 Δ*druE* mutant with pWKS30-encoded *druE* WT and the indicated *druE* variants. Data represent means of log_10_ EOP values ± SD of 3-4 biological replicates. Significance indicators were assessed by Dunnet’s 1-way-ANOVA multiple comparison tests. *, p ≤ 0.05; **, p ≤ 0.01; ***, p ≤ 0.001; ****, p ≤ 0.0001; ns, not significant. (D) FA assays monitoring binding of DruE^WT^ and the indicated truncation variants to 15-nt ssDNA. Data represent means ± SEM of three independent experiments using the same biochemical samples. (E) Intrinsic and DNA-stimulated ATPase activities of DruE^WT^ and the indicated truncation variants derived from thin-layer chromatographic analyses. Data represent means ± SEM of three independent experiments using the same biochemical samples. Significance was assessed *via* unpaired Student’s-t-tests. *, p ≤ 0.05; **, p ≤ 0.01; ns, not significant. (F) Unwinding of 3′ overhang (ovh) DNA by WT DruE and the indicated truncation variants monitored *via* stopped-flow/fluorescence assays. Single representative traces of three independent measurements are shown.

To explore the contributions of the identified domains to DruE nucleotide and DNA transactions, we created a series of truncations and characterized them *in vitro* and *in vivo* (**Figures 3A and S2**). The RecA1/2 core and inserted domains (DruE^1-1171^) lacked DNA affinity and ATPase activity, a fragment that also included the eWH and OBZ domains (DruE^1-1502^) bound ssDNA and exhibited DNA-stimulated ATP hydrolysis, while the MZB domain was additionally required for DNA unwinding (DruE^1-1723^; **Figure 3B-D**). The C-terminal PLDs were dispensable for all three functions (**Figures 3B-3D**), while DruE with a E525A substitution in RecA1 bound ssDNA but no longer hydrolyzed ATP or unwound dsDNA, as expected (**Figures S1C-S1E**). For *in vivo* testing of the functionality in anti-phage defense, we used the previously constructed *E. coli* EDL933 Δ*druE* deletion mutant, complemented with DruE variants produced from a plasmid.^26^ Using infection assays with phage Bas26 as the most sensitive Druantia target, we found that only one truncation (DruE^1-1723^) restored protection, while the other truncation variants as well as full-length DruE with the inactivating substitution in RecA1 (E525A) or residue exchanges in the ZBM of the MZB domain (C1681A/C1683A, C1687A/C1690A) failed to restore protection (**Figure 3E**). As DruE^1-1723^ lacks the C-terminal PLD domains, these results suggest that the PLDs are not involved in Druantia-mediated defense and instead could serve other roles beyond protection against Bas26.

### DruE dimerization is required for phage defense

In all five dimeric constellations, two DruE molecules consistently contact each other *via* their DDs in addition to more variable contacts between the DBDs (**Figures 4A and 4B**). The DD comprises four helices and dimerization is mediated by reciprocal interactions of the loops between the first and second helices and by an anti-parallel alignment of the third helices of the two protomers (**Figure 4C**). To determine whether the DD drives DruE dimerization, we generated and purified a DruE variant, in which the DD is replaced by a GSG linker (DruE^ΔDD^; **Figure S2**). Size-exclusion chromatography (SEC) in combination with multi-angle light scattering (MALS) detection revealed that DruE (calculated monomeric molecular mass = 237 kDa) eluted with an estimated molecular mass of 438 kDa, while DruE^ΔDD^ (calculated monomeric molecular mass = 229 kDa) eluted with an estimated molecular mass of 256 kDa (**Figure 4D**). The approximately twofold higher molecular mass observed for DruE, but not for DruE^ΔDD^, indicates that the DD is required for DruE dimerization.

**Figure 4.**
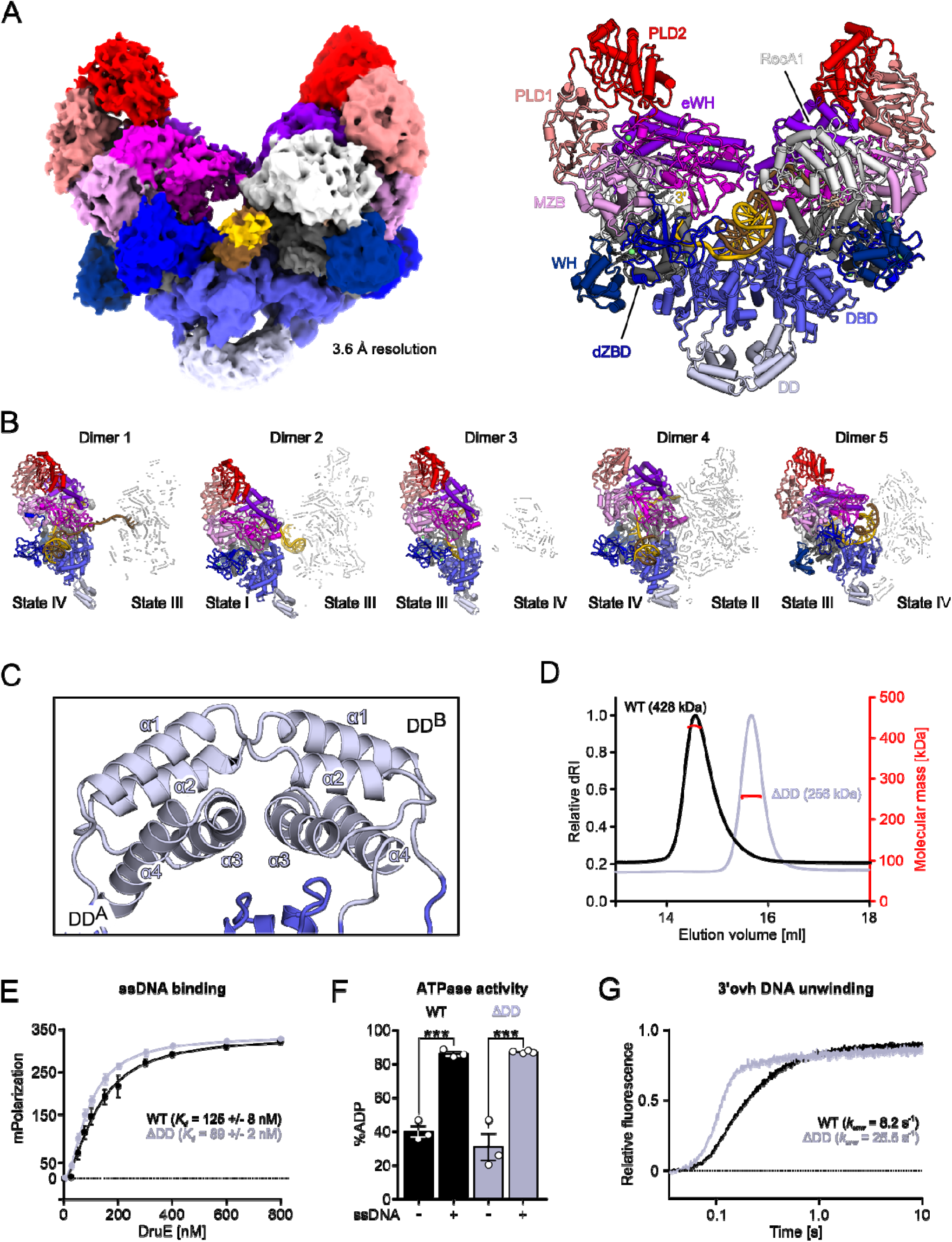
A small helical domain mediates dimerization of DruE required for phage defense. (A) CryoEM reconstruction (left) and cartoon representation (right) of a representative dimeric DruE (dimer 2) in complex with a forked DNA substrate and the weakly hydrolyzable ATP analog, ATPγS. (B) Cartoon representations of five dimeric structures of DruE in complex with a forked DNA substrate and the weakly hydrolyzable ATP analog, ATPγS. DruE protomers exhibit different conformational states (I-IV) as indicated. (C) Interaction of the DDs of the two DruE promoters of a representative DruE dimer (dimer 2). Secondary structure elements of the domains are labeled. (D) SEC-MALS analyses of DruE and DruE^ΔDD^. The relative differential refractive index (dRI) and the calculated molecular mass are plotted against the elution volume. (E) FA assays monitoring binding of DruE^WT^ and DruE^ΔDD^ to 15-nt ssDNA. Data represent means ± SEM of three independent experiments using the same biochemical samples. (F) Intrinsic and DNA-stimulated ATPase activities of DruE^WT^ and DruE^ΔDD^ derived from thin-layer chromatographic analyses. Data represent means ± SEM of 3-4 independent experiments using the same biochemical samples. Significance was assessed *via* unpaired Student’s-t-tests. *, p ≤ 0.05; **, p ≤ 0.01; ns, not significant. (F) Unwinding of 3′ overhang (ovh) DNA by WT DruE or DruE^ΔDD^ monitored via stopped-flow/fluorescence assays. Single representative traces of three independent measurements are shown.

We next asked whether DruE dimerization contributes to phage defense. Lack of the DD did not significantly affect the DNA affinity or DNA-stimulated ATPase activity of DruE *in vitro* (**Figures 4E and 4F**), and DruE^ΔDD^ unwound DNA about three times faster than DruE (**Figure 4G**). Despite the enhanced helicase activity, DruE^ΔDD^ failed to restore protection against Bas26 in the *E. coli* EDL933 Δ*druE* mutant strain (**Figure 3C**). These observations indicate that DruE dimerization facilitated by the DD is critical for full phage protection.

### DruE resorts to a unique molecular mechanism for DNA unwinding

Processive SF2 helicases unwind their substrates by threading a tracking nucleic-acid strand across the RecA domains, excluding the displaced strand, and undergoing ordered conformational changes driven by NTP binding, hydrolysis, release and re-binding to translocate into the duplex region one bp at a time. Notably, across our DruE structures, we observed individual protomers in four distinct DNA-bound states (I-IV), which correlate with distinct DruE conformations (**Figures 4B and 5A**).

**Figure 5.**
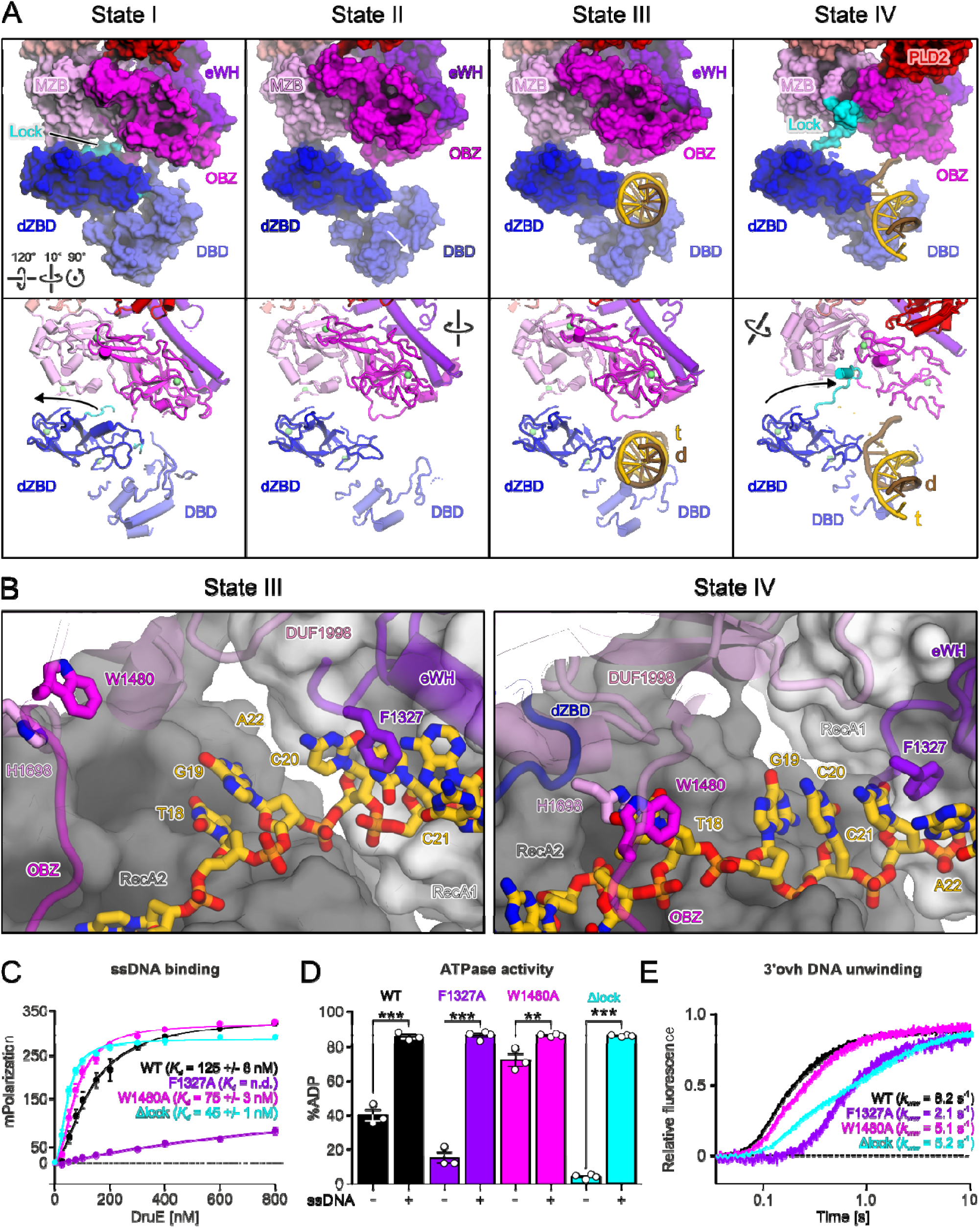
DruE adopts distinct conformational states and DNA contacts supported by unique molecular elements. (A) Surface (top) and cartoon (bottom) representations of DruE-ATPγS-DNA complexes in state I (dimer 2), state II (dimer 4), state III (dimer 5), and state IV (dimer 1), revealing major conformational rearrangements and a unique lock feature (cyan). For displayed protomers, see Figure 4B. (B) Changes in the interaction of DruE with the tracking strand upon transitioning from state III (dimer 5, left) to state IV (monomer, right). In state IV, the clamp element (residues W1480A and H1698) sandwiches a nucleobase of the tracking strand and the F1237A wedge intercalates between nucleobase stacks accommodated on the RecA domains. In state III, the corresponding elements are remote from the tracking strand. RecA domains are shown as surfaces, other domains as semi-transparent cartoons. Selected residues and nucleotides of the tracking strands are shown as sticks and colored by atom type; protein carbon, as the respective domains; RNA carbon, gold; nitrogen, blue; oxygen, red; phosphorus, orange. (C) FA assays monitoring binding of the indicated DruE variants to 15-nt ssDNA. Data represent means ± SEM of three or four independent experiments using the same biochemical samples. (D) Intrinsic and DNA-stimulated ATPase activities of the indicated DruE variants derived from thin-layer chromatographic analyses. Data represent means ± SEM of 3-4 independent experiments using the same biochemical samples. Significance was assessed *via* unpaired Student’s t-tests. *, p ≤ 0.05; **, p ≤ 0.01; ns, not significant. (E) Unwinding of 3′ overhang (ovh) DNA by the indicated DruE variants monitored *via* stopped-flow/fluorescence assays. Single representative traces of three independent measurements are shown.

In state I, the DBD is docked to the dZBD, and the OBZ and MZB domains close down on the dZBD and DBD domains like a lid (**Figure 5A**). A long loop of the dZBD (residues 505-523), which we term the “lock”, stabilizes the state I conformation by lining the RecA2, DBD, OBZ, and MZB domains. Only the canonical DNA-binding site of the RecA1 domain is partly accessible and bound by a portion of a DNA strand displaced from a neighboring DruE protomer. In state II, the DBD is displaced from the dZBD, leading to disengagement of the lock that is now disordered (**Figure 5A**). RecA, eWH, OBZ, MZB, and C-terminal domains rotate relative to the dZBD and DBD domains, rendering the canonical DNA-binding sites of RecA2 and neighboring surfaces of dZBD and DBD more accessible. In state III, the DNA tracking strand occupies binding sites unlocked in state II, running along the entire RecA platform, and the duplex portion of the DNA substrate comes to rest on top of the dZBD and DBD domains (**Figure 5A**). Finally, in state IV, the eWH/OBD/MZD/PLD region rotates relative to the dZBD/DBD duplex-binding platform; this rearrangement leads to (i) closing of the OBZ/MZB lid and repositioning the tracking strand across the RecA domains; (ii) deeper embedding of the duplex region on the dZBD/DBD duplex-binding platform; and (iii) formation of a DBD/OBZ channel, through which the displaced strand is now guided (**Figure 5A**). Notably, the dZBD lock in state IV spans across the tracking strand proximal to the fork junction and occupies a cleft between the OBZ and MZB domains, apparently stabilizing the observed conformation.

Irrespective of the conformational state, all DruE DNA-binding surfaces predominantly contact the DNA backbone, suggesting that the helicase binds and unwinds DNA sequence-independently. Supporting this notion, monomeric DruE and the state IV protomer of DruE dimer 1 (**Figure 4B**) exhibit 11- or 12-bp duplex regions, respectively, and a corresponding register shift of the tracking strand across its DNA-binding surfaces, also suggesting that initial full accommodation of a DNA substrate can lead to the unwinding of one or two bp.

While states I and II likely represent steps of initial DNA engagement and preparation for DNA accommodation, respectively, a more detailed comparison of states III and IV suggests that DruE toggles between these conformations during stepwise duplex unwinding. In state IV, the DNA duplex region rests in an “entry channel” formed by the dZBD and DBD domains (**Figure 5A**). The displaced strand is guided away from its complement through a channel formed between the DBD and OBZ domains, with the backbone running along the OBZ surface and the bases pointing towards the DBD. One ZBM in the OBZ domain engages regions in the displaced strand more distal from the duplex portion. The tracking strand runs across the DBD and between the dZBD and OBZ domains towards the RecA2 domain. W1480 (OBZ) and H1698 (MZB) form a “clamp” that sandwiches a nucleobase of the tracking strand (Thy18 in monomeric DruE), thereby guiding the following region of the 3′ overhang towards the RecA core. The tracking strand then traverses the RecA2 and RecA1 domains, engaging in contacts

to helicase motifs IV and V. The eWH, OBZ and MZB domains form a roof over the RecA domains, creating a DNA-binding tunnel through which the 3′ overhang is threaded. Consecutive 3-nt and 4-nt base-stacked blocks of the tracking strand (Cyt19-Ade21 and Ade22-Cyt26 in monomeric DruE) rest on the RecA2 and RecA1 domains, respectively, separated by F1327 (eWH) that wedges in between and extends the 4-nt stack. The very 3′ end of the tracking strand (Cyt27-Thy30 in monomeric DruE) turns away from the RecA1 platform and is anchored between the OBZ and eWH domains. In state III, in contrast, the OBZ/MZB lid atop the RecA domains remains open. As a consequence, the F1327 (eWH) wedge and the W1480 (OBZ)/H1698 (MZB) clamp are remote from the tracking strand and do not stack with the nucleobases as observed in state IV (compare **Figure 5B**). Furthermore, the DBD/OBZ channel for the displaced strand is closed and the dZBD lock is disordered (**Figure 5A**). Morphing between states III and IV illustrates how insertion of the F1327 wedge into the tracking strand and engagement of the W1480/H1698 clamp could be correlated with translocation of DruE into the duplex by one bp (**Movie S1**). Overall, states I-IV capture a series of discrete events, from initial binding of the DNA substrate by the RecA1 domain to full engagement aided by DruE-specific auxiliary domains that accommodate the duplex region and help direct the tracking and displaced strands.

### The distinct helicase mechanism of DruE is required for phage defense

Based on our structural analysis, the dZBD lock, F1327 wedge, and W1480/H1698 clamp are DruE-specific elements involved in DNA unwinding. To test the functional importance of these elements, we created DruE variants that lacked residues 510-526 (DruE^Δlock^) replaced F1327 or W1480 with alanine (DruE^F1327A^, DruE^W1480A^) and tested these variants for DNA binding, ATPase, and helicase activities.

Compared to DruE, DruE^Δlock^ exhibited 2.7-fold higher affinity to ssDNA (**Figure 5C**). Furthermore, while the DNA-stimulated ATPase activity of DruE^Δlock^ was largely unaffected, DruE^Δlock^ showed a tenfold lower intrinsic ATPase activity than DruE (**Figure 5E**). These observations are consistent with the lock adopting a DNA-competitive conformation in state I (compare **Figure 5A**), mimicking a bound tracking strand and thereby leading to increased intrinsic ATPase activity. DruE^Δlock^ (*k* = 5.2/s) also showed a 40 % reduction in DNA unwinding activity compared to DruE (*k_unw_* = 8.2/s; **Figure 5D**). Furthermore, plasmid-based expression of *druE*^Δlock^ in the EDL933 Δ*druE* strain failed to restore phage defense against Bas26 (**Figure 3C**). As the lock adopts different conformations in states III and IV, DruE conformational toggling is likely required for DNA unwinding, while the lock-supported helicase activity is essential for phage defense.

F1327 is repositioned by about 8 Å in state III compared to state IV, resulting in a disruption of F1327-nucleobase π-stacking in state III (**Figure 5B**). Consistent with a role in intermittent DNA binding, the DNA affinity of DruE^F1327A^ was strongly reduced compared to the wildtype (WT; **Figure 5C**). While, nevertheless, the DNA-stimulated ATPase activity was largely unchanged (**Figure 5D**), DNA unwinding of DruE^F1327A^ was strongly impaired (about 30% of WT DruE; **Figure 5E**). These results underscore an important role of F1327 during DruE-mediated DNA unwinding. By switching between released (state III) and a nucleobase-stacked (state IV) constellations, F1327 may function as a ratchet, as previously proposed for a long α-helix in Ski2-like helicases^38^. Furthermore, efficiency of plaquing (EOP) assays showed that ectopically expressed pWKS30-*druE*^F1327A^ in the EDL933 Δ*druE* mutant did not restore the phage defense against Bas26 to the WT level (**Figure 3C**), indicating that F1327-dependent DruE helicase activity is required for phage defense.

Similar to F1327, the W1480/H1698 clamp switches between unbound (state III) and DNA-bound (state IV) conformations (**Figure 5B**). While DruE^W1480A^ showed slightly enhanced DNA affinity and increased intrinsic ATPase activity compared to WT DruE (**Figure 5D**), it unwound DNA at a comparable rate (**Figure 5E**). Still, complementation of plasmid-expressed *druE*^W1480A^ in the Δ*druE* mutant only partially restored phage defense against Bas26 (**Figure 3C**), suggesting that the W1480/H1698 clamp is important for DruE helicase activity, but that H1698 alone can partially maintain DNA-binding as observed in state IV. Taken together, these findings indicate that DruE resorts to a hitherto unobserved mode of DNA unwinding that relies on lock, wedge, and clamp motifs in DruE-specific auxiliary domains.

### DruE interacts with DruH and other host proteins under non-infecting conditions

To obtain insights into the molecular functions of DruH, we purified the protein from *E. coli* (**Figure S2A**) and elucidated its structure by cryoEM (**Figures S9 and S10**; **Table S1**). DruH adopts an oval-shaped structure with a circular arrangement of ten domains designated D1 to D10. (**Figures 6A and 6B**). Searches for related folds *via* the DALI server^39^ did not reveal any known catalytic domains in DruH. However, three domains (D6, D7, D8) resemble the fold and topology of Fn3-like domains (**Figure 6C**). The Fn3 fold comprises seven β-strands arranged in two sheets with four and three strands, respectively. Fn3-like domains are part of the immunoglobulin (Ig) superfamily and occur in a large variety of proteins, such as cell surface proteins, cytokine receptors, and chaperonins^40^. However, we could not discern conserved sequence motifs, such as the cell surface-binding RGD motif^41^, in these three domains. D4 adopts the related intermediate Ig-like fold ^40^ (**Figure 6C**), which also pertains to the Ig superfamily and merely differs from Fn3-like domains in the position of one β-strand^41^. As Ig-like domains frequently mediate protein-protein interactions, their presence in DruH suggests that DruH might act as a scaffold for other proteins associated with the Druantia III-A phage defense mechanism or for phage recognition. However, DruH did not stably interact with purified DruE in SEC (**Figure 6D**), irrespective of the presence of DNA or various nucleotides (**Figure S11A**). Furthermore, we did not observe modulation of the DruE helicase activities by DruH *in vitro* (**Figure S11C**).

**Figure 6.**
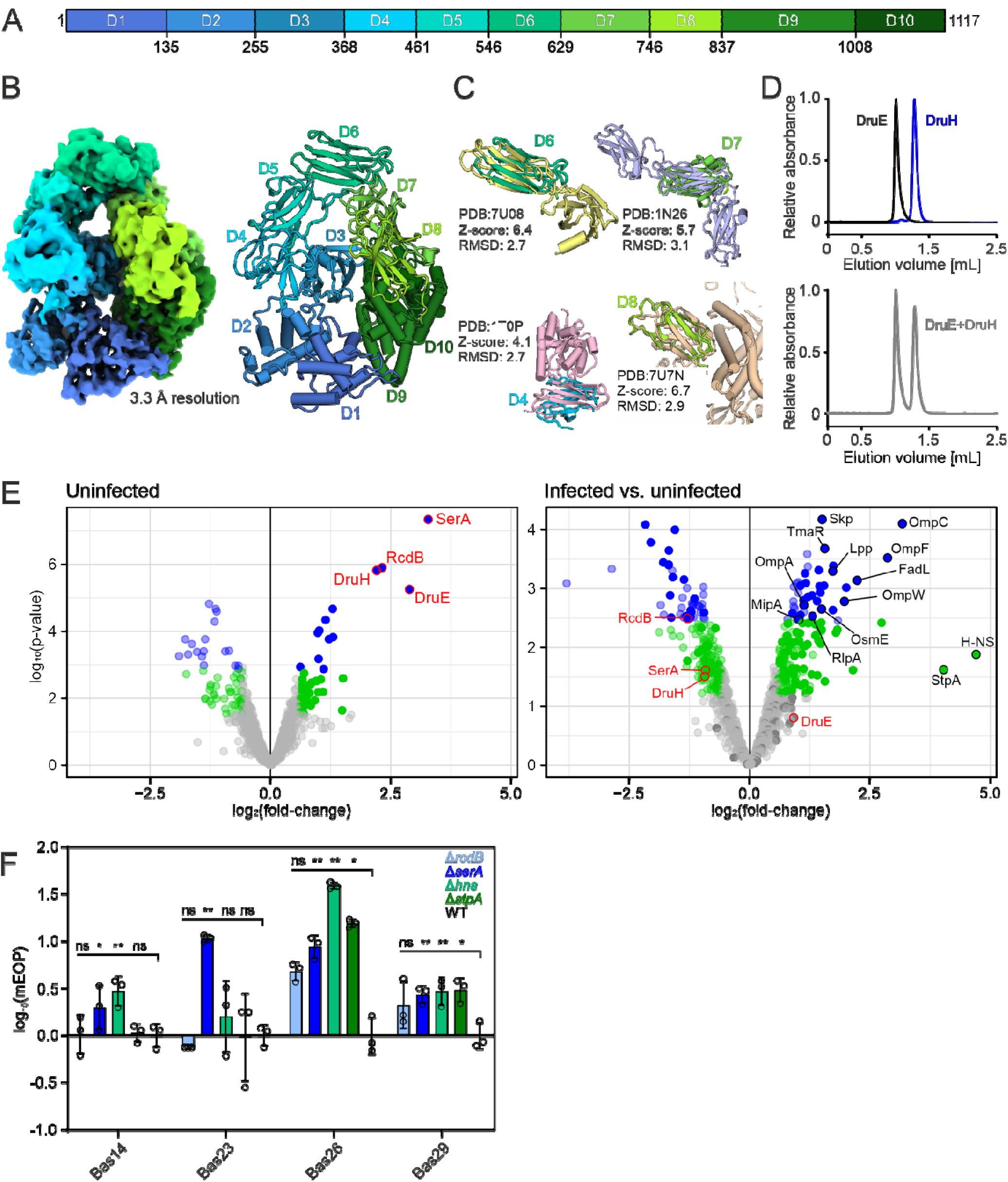
DruH indirectly interacts with DruE *in vivo* alongside additional host proteins released upon infection. (A) Domain organization of DruH. DruH can be divided into ten domains denoted D1 - D10. (B) CryoEM reconstruction (left) and cartoon representation (right) of DruH colored by domains as in (A). (C) Superposition of the indicated DruH domains with the closest structural homologs identified by a DALI search^39^. DruH domains D6, D7, and D8 resemble the Fn3-like domains of CD148 (PDB-ID: 7U08), the human interleukin-6 receptor alpha-chain (PDB-ID: 1N26), and IL-27 (PDB-ID: 7U7N), respectively. D4 adopts an intermediate Ig-like fold also found in the ICAM-3 N-terminal domain (PDB-ID: 1T0P). (D) UV_280_ traces of analytical SEC runs of DruE and DruH analyzed separately (upper chromatogram) and after mixing (lower chromatogram). (E) Volcano plots of proteins enriched following DruE pulldown under infected or uninfected conditions as determined by mass spectrometry. Infections were conducted with Bas23 phage. Left: protein enrichment when pulling down DruE with a C-terminal 3xFLAG tag versus an untagged control. Right: relative protein enrichment comparing infected conditions to uninfected conditions. Grey dots, no-hit proteins. Green dots, candidate proteins. Blue dots; enriched hits. Red outline, selected enriched hits under uninfected conditions. Black outline, enriched or candidate hits involved in membrane synthesis, membrane transport, membrane protein organization, or DNA binding. Values represent the mean of three biological replicates. (F) Relative contribution of identified enriched proteins to anti-phage defense by Druantia-III in *E. coli* ATCC8739. mEOP, or modified efficiency of plating divides the EOP of the single Δ*serA*/*rcdB*/*hns*/*stpA* deletion mutants by the EOP of the Δ*druHE* Δ*serA*/*rcdB*/*hns*/*stpA* double-deletion mutants. EOP is calculated based on plaque formation of the background strain lacking the tested mutation (i.e., WT for each single deletion, Δ*druHE* for eacg double deletion). Bars and error bars represent the mean ± SD from three biological replicates. Open circles represent values from individual experiments. Significance was assessed *via* Benjamini, Krieger, and Yekutieli multiple unpaired t-tests. *, q ≤ 0.05; **, q ≤ 0.01; ***, q ≤ 0.001; ns, not significant.

Despite the absence of detectable binding between DruE and DruH *in vitro*, we examined whether the two proteins interact *in vivo*, potentially together with additional binding partners. We initially designed co-immunoprecipitation experiments followed by western blotting analysis between DruE and DruH. However, all tested tag types and tag positions on DruH resulted in a loss of the defense phenotype (**Figure S12**). In contrast, tagging DruE with a 3xFLAG tag at either the N- or C-terminus did not interfere with phage defense (**Figure S12**). We therefore tagged DruE at the C-terminus in the plasmid with the native Druantia III-A system from *E. coli* ATCC8739. To identify protein partners in the native context, we introduced the plasmid with tagged DruE into the *E. coli* ATCC8739 strain with *druHE* deleted, co-immunoprecipitated proteins under non-infecting conditions or when infected with Bas23, and analyzed pulled-down proteins by mass spectrometry (**Table S2**).

Under uninfected conditions, DruH, the D-3-phosphoglycerate dehydrogenase SerA, and the transcriptional regulator RcdB were highly enriched compared to the untagged control (**Figure 6E, left**). Under infecting conditions, a large number of enriched hits were proteins associated with bacterial membrane organization and pore-forming proteins known to act as common receptor-binding proteins (RBPs) for phages (**Figure 6E, right**). Comparing the change in enrichment between non-infecting and infecting conditions revealed relative depletion of DruH, SerA, and RcdB, indicating that their interaction with DruE is lost as part of an immune response (**Figure 6E, right**). We also observed an enrichment of the histone-like proteins H-NS and StpA (**Figure 6E, right**), which form heterodimers, are known to interfere with phage infections by binding AT-rich DNA, and are inhibited by some phage-encoded factors^42–44^. Notably, phages that are not sensitive to heterologous expression of Druantia III-A (**Figure 1B**), such as members of the *Autographivirinae* and *Tevenvirinae* families, encode anti-H-NS proteins, including MotB and Arn in phage T4^45–47^. Overall, these results show that DruE interacts with DruH and other host proteins that dissociate during an immune response.

We finally asked if the host proteins interacting with DruE before or during an infection contribute to Druantia’s defense. We specifically focused on SerA and RcdB as the two most enriched host proteins under non-infected conditions as well as H-NS and StpA given their binding of phage DNA. We generated knockouts of each gene in *E. coli* ATCC8739 with the endogenous Druantia III-A system intact or deleted and measured infectivity by four phages countered by Druantia III-A (Bas14, Bas23, Bas26, and Bas29). To determine the extent to which each gene contributes to Druantia’s defense, we compared the enhanced protection for each gene with or without Druantia III-A. While none of the genes were essential for Druantia’s defense (*i.e.*, modified [m] EOP = 100), each significantly contributed to the system’s defense for at least two phages and *serA* and *hns* for all four tested phages (**Figures 6F and S13**). Taken together, these results show that the Druantia III-A system involves other host proteins involved in phage defense and other seemingly unrelated cellular processes to fend off foreign invaders.

## DISCUSSION

Based on our findings, we propose a working model of anti-phage defense by Druantia III-A systems (**Figure 7**). Upon infection, a DruE dimer in complex with DruH and other host proteins, such as SerA and RcdB, engages exposed ssDNA encountered in the bacterial cytosol. The engaged DNA can be unmodified or methylated, while other chemical modifications block immunity. Engagement of both DNA strands by the DruE dimer presumably initiates the immune response, which involves ATP-dependent 3′-to-5′ DNA unwinding by DruE through conformational shifts resolved in our cryoEM structures, resulting in the displacement of the interacting host proteins in favor of a new contributing interactome. Defense depends on a unique unwinding mechanism, implemented *via* distinct lock, wedge, and clamp molecular tools that are supplied by DruE-specific auxiliary domains. This process leads to clearance of the recognized invader, albeit through currently unknown mechanisms.

**Figure 7.**
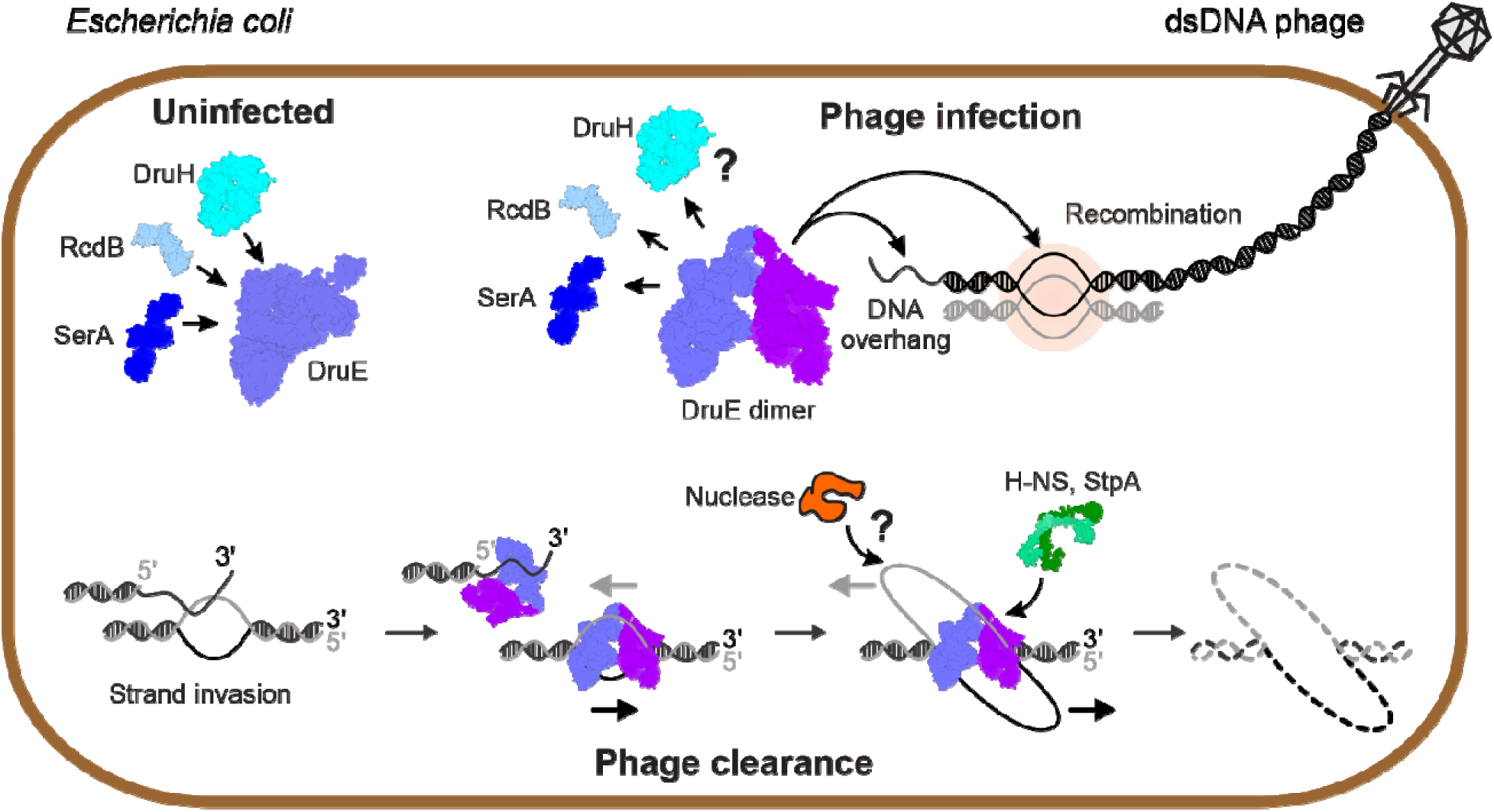
Working model for phage clearance by Druantia III-A. Druantia III-A relies on both system components, DruH and DruE, which interact in the absence of an infection and are partly mediated by the host proteins SerA and RcdB. Upon infection, the interaction partners of DruE are displaced. Regions of single-stranded DNA present in the invading and replicating phage DNA are engaged by DruE, which subsequently unwinds neighboring duplex regions to initiate phage clearance. Clearance is assisted by additional host-factors, such as H-NS and StpA.

Apart from the mechanistic basis of phage clearance, additional aspects of the immune system warrant further investigation. First, the exposed ssDNA that triggers the Druantia III-A system could come in different forms, such as ssDNA ends in the injected phage DNA or as intermediate of recombination inherent to DNA replication in some phages, including Bas29 and Bas54^33,48–50^, shown to be sensitive to Druantia in our work (**Figure 1B**). The conversion of Druantia III-A to a cytotoxic system through the addition of mitomycin C suggests that Holliday junctions or other intermediates that occur during recombination are preferentially recognized, although the same recognition could pose a real risk to cells facing harsh environmental conditions – a hazard also posed by Hachiman that is triggered by chromosomal DNA damage^51^.

Another aspect deserving further investigation is whether DNA cleavage plays a role in defense by Druantia III-A. DruE has been proposed to possess nuclease activity through its PLD2 that would lead to invader clearance through DNA cleavage^25^. However, we did not observe any reproducible nuclease activity *in vitro* (**Figure S3B**), and also in the varying structures derived from cryoEM no cleavage of ssDNA portions was noted. Additionally, the PLDs were fully dispensable for phage defense. PLDs have also been associated with membrane association^52^, although TIRF microscopy showed that a functional DruE-mNeonGreen fusion displayed cytosolic localization under infecting and non-infecting conditions (**Figure S14**). We instead speculate that the PLDs play an important role outside of our specific experiments, such as mediating apparent synergy between Druantia III-A and co-encoded Zorya II and ARMADA II systems^25,26^ not explicitly tested in this work. DruE had also been connected to flagellar biosynthesis and motility in *E. coli* EDL933^53^, although we found no such effect of deleting *druE* in this strain based on Northern blotting analysis of *fliC* or on motility assays (**Figure S15**).

DruH also remains enigmatic, although our findings provide key insights helping to inform its role in immunity. Despite DruH lacking any identifiable catalytic domains, our cryoEM structure revealed a monomer with multiple Ig-like domains known for forming protein-protein interactions^40^. Given that DruH interacts with DruE only in a cellular context with other host proteins and is displaced upon immune activation, DruH could offer a non-catalytic protein scaffold or could help discern ssDNA associated with phages that kickstarts immunity. The identification of the D-3-phosphoglycerate dehydrogenase SerA involved in L-serine biosynthesis^54^ and the transcriptional regulator RcdB involved in biofilm formation^55^ as part of the interactome of DruE and DruH offered some support for a scaffold-like function of DruH, although the involvement of these host proteins and their contribution to Druantia-mediated phage defense remains mysterious given that neither protein plays any obvious role in anti-phage defense.

Despite these open questions, our work sheds light on the properties of other Druantia types and subtypes as well as other known or putative anti-phage defenses with a common YprA/MrfA-like helicase that remain uncharacterized. The III-B subtype, which also comprises only DruE and DruH, possesses a more compact DruE lacking the PLDs that were dispensable in our experiments with the III-A system^25^. The identification of seemingly unrelated host proteins as contributing binding partners would suggest that the III-B systems and possibly other Druantia types could benefit from host factors. The additional genes within the larger Druantia types could substitute for these host factors, although none share notable homology or annotated domains with the identified binders^24^.

Beyond the various Druantia types and subtypes, the immune systems are linked to a recently expanded set of defenses united by a YprA/MrfA-like helicase distinguished by the MZB domain (DUF1998) at varying distance C-terminal of the RecA core^25,27,56^. Such helicases are also found in the ARMADA, DISARM, Dpd, BRIGADE, and TALON phage defense systems^25^. Of these systems, DISARM is the best characterized to date, reported to restrict incoming phage DNA and methylate host DNA, reminiscent of restriction-modification systems^24^. The DISARM system relies on the YprA/MrfA helicase DrmAB that recognizes dsDNA with a 5′ overhang ^20^, however the underlying phage clearance mechanism remains elusive. While DISARM also relies on the recognition of ssDNA, the ability of Druantia III-A to recognize any ssDNA suggests that these two defenses, and possibly the other defenses in the YprA/MrfA-like helicase family, recognize distinct forms of ssDNA that trigger defense. In addition, DISARM was reported to have the opposite directionality of DNA unwinding (5′-to-3′) and relies on a “trigger loop” that regulates substrate specificity^20^, as opposed to the dZDB lock of DruE, which supports duplex unwinding. DISARM also contains PLDs in a separate protein, DrmC, that was dispensable for defense against some phages and is not found in all DISARM systems^25^, further suggesting that these common (but not universal) domains in the YprA/MrfA-like helicase family play a yet-to-be revealed role. Thus, our work suggests that more functional diversity awaits discovery through further characterization of the Druantia types and subtypes as well the YprA/MrfA-like helicase family of defenses.

## RESOURCE AVAILABILITY

### Lead contact

Requests for further information and resources should be directed to and will be fulfilled by the lead contact, Chase Beisel.

### Materials availability

All unique/stable reagents generated in this study are available from the lead contact with a completed materials transfer agreement.

### Data and code availability

Raw EM data are available from the lead contact upon request. CryoEM reconstructions have been deposited in the Electron Microscopy Data Bank (https://www.ebi.ac.uk/pdbe/emdb) under accession codes EMD-56257 (https://www.ebi.ac.uk/pdbe/entry/emdb/EMD-56257; DruH), EMD-56251 (https://www.ebi.ac.uk/pdbe/entry/emdb/EMD-56251; DruE monomer), EMD-56252 (https://www.ebi.ac.uk/pdbe/entry/emdb/EMD-56252; DruE dimer 1), EMD-56253 (https://www.ebi.ac.uk/pdbe/entry/emdb/EMD-56253; DruE dimer 2) EMD-56254 (https://www.ebi.ac.uk/pdbe/entry/emdb/EMD-56254; DruE dimer 3), EMD-56255 (https://www.ebi.ac.uk/pdbe/entry/emdb/EMD-56255; DruE dimer 4), and EMD-56256 (https://www.ebi.ac.uk/pdbe/entry/emdb/ EMD-56256; DruE dimer 5). Structure coordinates have been deposited in the RCSB Protein Data Bank (https://www.rcsb.org) with accession codes 9TUD (https://www.rcsb.org/structure/9TUD; DruH) 9TU7 (https://www.rcsb.org/structure/9TU7; DruE monomer), 9TU8 (https://www.rcsb.org/structure/9TU8; dimer 1), 9TU9 (https://www.rcsb.org/structure/9TU9; dimer 2), 9TUA (https://www.rcsb.org/structure/9TUA; dimer 3), 9TUB (https://www.rcsb.org/structure/9TUB; dimer 4), and 9TUC (https://www.rcsb.org/structure/9TUC; dimer 5). The full Mascot search results of DruE-mNeonGreen and LacZ are available at https://box.fu-berlin.de/s/4Sp5GKFBscALqwx.

Mass spectrometry raw data will be accessible on the PRIDE database.

This paper does not report original code.

Any additional information required to reanalyze the data reported in this paper is available from the lead contact upon request.

## ACKNOWLEDGMENTS

We thank Nicole Holton, Laboratory of Structural Biochemistry, Freie Universität Berlin, Eva Absmeier, Laboratory of mRNA Translation and Turnover, Freie Universität Berlin, and Luise Franz, Leibniz-Foschungsinstitut für Molekulare Pharmakologie, Berlin, for help with molecular cloning and initial protein production and characterization, as well as Nikos Arkoulakis, Jan-Hendrik Latz, and Leonie Nowara, Institute of Biology-Microbiology, Freie Universität Berlin, for initial growth assays of the Δ*druE* mutant and complemented strains. This work was supported by grants from the Deutsche Forschungsgemeinschaft (DFG) in the framework of the priority program SPP 2330 (project number 548567920 to P.F.P [PO 2831/2-1] and M.E. [ER 778/13-1]; project number 465069819 to C.L.B. [BE 6703/2-2]) and in the framework of the research training group RTG 2473 (project number 392923329 to M.C.W.). We acknowledge the assistance of the core facility BioSupraMol supported by the Deutsche Forschungsgemeinschaft in electron microscopic and mass spectrometric analyses.

## AUTHOR CONTRIBUTIONS

S.H., T.G., L.M.G., H.L., V.V.L., C.C., and E.K. conducted all experiments, except TIRF microscopy, mass spectrometric analysis and cryoEM data collection and processing. P.F.P. performed TIRF microscopy. B.K. conducted mass spectrometric analyses. T.H. acquired and processed cryoEM data. L.M.G., S.H., and B.L. built atomic models and refined structures. S.H., T.G., L.M.G., C.L.B., and M.C.W. wrote the manuscript. All authors participated in data analysis and interpretation. M.E., H.A., C.L.B., and M.C.W. conceived the project, supervised the work in their laboratories and coordinated collaborations.

## METHODS

### Molecular cloning and protein production

Oligonucleotides used in this study are listed in **Table S3**. Bacterial strains and plasmids used in this study are listed in **Table S4**.

To generate the pDruHE vector, the Druantia III-A operon was PCR-amplified from the *E. coli* ATCC8739 genome in two fragments and inserted into the linearized p15a_empty vector by reverse PCR. PCR fragment sizes were verified by electrophoresis on an agarose gel with Midori Green, purified with the PCR clean and concentrator kit (Zymo Research), assembled with NEBuilder HiFi DNA assembly Gibson reagents (NEB) and transformed into chemically competent Top10 *E. coli* by a heat shock at 42°C for 30 s. Cells were transferred into SOC medium at 37°C for 1 h, and then plated on a chloramphenicol-containing agar plate. pDruH and pDruE were generated by reverse PCR on pDruHE and circularized with KLD mix (NEB). For the complementation assays, the chloramphenicol resistance cassette was swapped with an ampicillin resistance cassette by reverse PCR and Gibson cloning on pDruH. pDruHE-3F was generated by adding a region encoding a triple flag tag downstream the *druE* gene of the pDruHE plasmid by Gibson cloning. All the mutant versions of *druE* were generated pDruHE by PCR-fragmentation into three pieces. 80-nt oligos carrying the desired mutation were used in a Bridging Gibson strategy with the other p15a_DTIII-A fragments (0.1 pmol of vector fragments, 0.4 pmol of oligos). All sequences were verified by Nanopore sequencing (Plasmidsaurus).

Electrocompetent *E. coli* MG1655, *E. coli* ATCC8739 and *E. coli* ATCC8739 Δ*druHE* cells were transformed with the different vectors (1.8 kV; 25 µF; 200 Ω; 1 mm cuvette), recovered as described above, and plated on chloramphenicol plates. Those strains were used for plaque assays, liquid culture, mitomycin C survivability, and pulldown experiment.

Synthetic gene blocks for a codon-optimized DruE coding region were stepwise assembled and integrated into a pFL vector using Gibson assembly ^57^ for production of DruE including a TEV-cleavable N-terminal His_10_-tag. Plasmids with DNA fragments encoding DruE variants were PCR amplified from the pFL-*druE* vector *via* inverse PCR^58^. In DruE^ΔDD^ the DD (residues 678-759) were replaced by a GSG linker. For DruE^Δlock^ the lock (residues 510-526) were replaced by a GSG linker.

The pFL constructs were transformed into *Escherichia coli* DH10 MultiBac cells, and blue-white screening was used to select colonies with successful Tn7 transposition into the baculovirus genome^59^. Bacmid DNA was purified and used to transfect SF9 insect cells with X-tremeGENE^TM^ gene 9 DNA transfection reagent (Roche Applied Science). The first virus generation, V_0_, was harvested and used to infect HighFive insect cells to produce the second virus generation, V_1_. The V_1_ virus was used to infect HighFive cells for protein production. Cells were harvested before the cell viability decreased below 92%.

A codon-optimized version of the *druH* gene was inserted into a pETM11 vector using *Nco*I and *Hind*III restriction sites for production of DruH with an N-terminal, TEV-cleavable His_6_-tag. pETM11-*druH* was transformed into *E. coli* Star (DE3) cells (Invitrogen). For protein production, cells were cultivated in auto-inducing medium^60^ at 37°C, 800 rpm, to an OD_600_ of 0.8, and for another 20 h at 18°C, 800 rpm, before harvest.

### Construction of EDL933 Δ*druE* complemented and ATCC8739 Δ*serA,* Δ*rcdB,* Δ*hns*, and Δ*stpA* strains

The EHEC O157:H7 Δ*stx1/2* strain EDL933 was used as native host for construction of complemented strains, expressing WT *druE*, the *druE*-mNeonGreen translational fusion and *druE* variants in the Δ*druE* deletion mutant^26^. The EDL933 Δ*stx1/2* Δ*druE* scarless deletion mutant was previously constructed using the one-step-inactivation method^26,61^. To complement the Δ*druE* mutant with WT *druE* and variants, the pFL constructs encoding DruE variants, which were codon-optimized for insect cell expression, were amplified by PCR and cloned into the pWKS30 plasmid^62^ by Gibson assembly. For complementation of the Δ*druE* mutant with *druE*-mNeonGreen, *druE* was amplified from strain EDL933 Δ*stx1/2* and fused in frame with mNeonGreen, amplified from plasmid pEM8731^63^ and cloned into pWKS30 using Gibson assembly.

The Δ*serA,* Δ*rcdB,* Δ*hns*, and Δ*stpA* mutants of *E. coli* strain ATCC8739 were constructed using the λ Red recombineering method^61^. Target operons were deleted from the first start codon to the last stop codon. A kanamycin resistance (*kanR*) cassette was amplified by PCR using Q5 High-Fidelity DNA Polymerase (NEB), with primers containing 50- to 60-bp homology arms corresponding to the flanking regions of the operons. Electrocompetent cells were prepared from strains harboring the temperature-sensitive pKD46 plasmid. Cultures were grown in LB supplemented with ampicillin (100 μg/ml) at 30°C. Production of recombination machinery was induced by adding 0.2% L-arabinose at an OD_600_ between 0.3 and 0.4. Cultures were harvested at OD_600_ = 0.6, washed with ice-cold 10% glycerol, and electroporated with 1,000 ng of purified PCR product (kanR cassette). After electroporation, cells were recovered in SOC at 37°C for 4 h to allow recombination and pKD46 plasmid curing. Cells were then plated on kanamycin (50 μg/ml) plates and incubated overnight at 37°C. Loss of pKD46 was confirmed based on ampicillin sensitivity. To remove the *kanR* cassette, verified mutant strains were transformed with the FLP recombinase-expressing plasmid pCP20. Electroporated cells were recovered in SOC at 30°C for 1 h, plated on LB agar with ampicillin (100 μg/ml), and incubated overnight at 30°C. Single colonies were re-streaked onto antibiotic-free LB agar and incubated at 40°C overnight to induce FLP recombinase activity and eliminate the pCP20 plasmid. Clones were tested for antibiotic sensitivity to both kanamycin and ampicillin, and verified by colony PCR and Sanger sequencing.

### Protein purification

Cell pellets were resuspended in 50 mM Tris-HCl, pH 8.0, 500 mM NaCl, 15 mM imidazole, 2 mM β-mercaptoethanol. Bacterial cell resuspensions were supplemented with 0.02% (v/v) NP40, 0.2 mg/ml lysozyme, and 0.01 mg/ml DNase. Insect cell resuspensions were supplemented with 0.02% (v/v) NP40, 0.01 mg/ml DNAse, 0.07 mg/ml AEBSF, and 0.1 mg/ml benzamidine hydrochloride. The cell suspension was lysed by sonication using a Sonopuls HD 3100 ultrasonic homogenizer (Bandelin). Cleared lysate was passed through 2 ml of Ni^2+^-NTA agarose beads (Qiagen) in a gravity flow column to capture the His_6/10_-tagged proteins of interest. After washing and elution with 300 mM imidazole, the N-terminal His_6/10_-tag was cleaved by incubation with TEV protease overnight. The samples were concentrated and further purified by size exclusion chromatography on a 26/60 Superdex 200 or 16/60 Superdex 200 column (Cytiva) in 10 mM Tris-HCl, pH 8.0, 200 mM NaCl, 1 mM DTT.

### Nucleic acid binding analyses *via* fluorescence anisotropy

5 nM 5-FAM-labeled ssRNA or ssDNA were titrated with increasing concentrations of DruE (WT or variants) in 40 mM Tris-HCl, pH 7.5, 150 mM NaCl, 5 mM MgCl_2_, 1 mM DTT, 46 mU/μl RNasin (for RNA samples). Experiments conducted with DruH were performed at lower salt concentration in 40 mM Tris-HCl, pH 7.5, 75 mM NaCl, 5 mM MgCl_2_, 1 mM DTT to enhance binding. Fluorescence anisotropy was quantified in 384-well plates using a Spark Multimode Microplate Reader (Tecan). To extract binding constants, changes in anisotropy were plotted against protein concentration, and the data were fitted to a single exponential Hill function (fraction bound = *A*[protein]^n^/([protein]^n^+*K* ^n^); *A*, fitted maximum of nucleic acid bound; n, Hill coefficient; *K_d_*, dissociation constant)^64^ using OriginPro (OriginLab 2025).

### Electrophoretic mobility shift assays

For EMSAs, we employed a chemically synthesized, 5′ Cy5-labeled 60-nt ssDNA probe (IDT) or dsDNA generated by annealing the ssDNA probe to a chemically synthesized complementary strand at a molar ratio of 1:50 and PAGE purification using the ZR small-RNA PAGE Recovery Kit (Zymo Research). DNA probes were incubated or not with recombinant DruE or DruE^1-1171^ at a ratio of 1:100 for 1 h at room temperature in 20 mM NaH_2_PO_4_, pH 8.0, 500 mM NaCl, 10 % (w/v) sucrose. 10% (v/v) glycerol were added to the samples before electrophoresis on a native TBE polyacrylamide gel (10%; 29:1) for 2 h at 75 mV and 4°C. Gels were imaged on an ImageQuant 800 imaging system (Amersham; Cy5 excitation wavelength; 30 s exposure).

### ATPase assays

DruE variants were diluted to 2.5 μM in 40 mM Tris-HCl, pH 7.5, 150 mM NaCl, 1 mM DTT, without or with 2 μM nucleic acid (ssDNA or ssRNA). The reactions were initiated by the addition of 1 mM ATP/MgCl_2_, supplemented with 0.0125 mCi/ml [α-^32^P]-ATP, and samples were incubated for 30 min at 25°C. The reactions were stopped by the addition of an equivalent volume of 100 mM EDTA, pH 8.0, and separated by thin-layer chromatography in 20% (v/v) ethanol, 6% (v/v) acetic acid, 0.5 M LiCl. Chromatograms were scanned on a Storm 860 phosphorimager (GMI) and quantified by densitometry.

### Helicase assays

RNA or DNA unwinding was monitored by fluorescence stopped-flow measurements on an SX-20MV spectrometer (Applied Photophysics). Nucleic acid substrates contained the same 12-base pair duplex regions and 31-nt single-stranded 3′ or 5′ overhangs (**Table S3**). The short strand was labeled with Alexa Fluor 488 at the 3′ end and the long strand, bearing the overhang, was labeled with the quencher Atto 540 Q at the 5′ end (in the case of the blocked overhang construct a third unlabeled strand was added), such that fluorophore and quencher resided close to each other on the blunt ends of the duplex regions after strand annealing. The fluorescence of Alexa Fluor 488 was excited at 465 nm and the fluorescence emission was monitored after passing a 495 nm cut-off filter (KV 495, Schott). An increase in fluorescence was only observed when the Alexa Fluor 488-labeled strand was separated from the complementary strand carrying the quencher Atto 540 Q. For all experiments, 1 µM DruE (WT or variants) was mixed with 50 nM RNA or DNA in 40 mM Tris-HCl, pH 7.5, 150 mM NaCl, 0.5 mM MgCl_2_. Experiments conducted with DruH as well DruE and DruH were also performed with 1 µM protein concentration but using 40 mM Tris-HCl, pH 7.5, 75 mM NaCl, 0.5 mM MgCl_2_ to enhance differences in unwinding activity. Mixtures were loaded into the syringe and incubated for 5 min at 25°C. 60 µl of the mixtures were then rapidly mixed with 60 µl of 4 mM ATP/MgCl_2_ in the same buffer and the fluorescence change was monitored over time.

Data from fluorescence measurement were baseline-corrected by subtracting the starting fluorescence immediately after addition of ATP and normalized to the baseline-corrected maximum fluorescence. Fluorescence traces for DruE/3′ overhang RNA, DruE/5′ overhang DNA, DruE^1-1502^/3′ overhang DNA, DruE^E252A^/3′ overhang DNA and DruE^1570-2104^/3′ overhang DNA, which did not show any unwinding, were normalized to the baseline-corrected maximum fluorescence of DruE/3′ overhang DNA. Three to five traces were averaged and plotted using Prism software (GraphPad). The data were fitted to a double exponential equation (fraction unwound = *A_fas_*_t_·(1-exp(-*k_fast_t*))+*A_slow_*·(1-exp(-*k_slow_t*)); *A_fast/slow_*, unwinding amplitude of the fast/slow phase; *k_fast/slow_*, unwinding rate constants of the fast/slow phase [s^-1^]; t, time [s]). The first s of data acquisition was excluded from curve fitting to account for the initial mixing periods. Amplitude-weighted unwinding rate constants were calculated as *k_unw_*=∑(*A_i_k_i_*^2^)/∑(*k_i_A_i_*).

### Nuclease assays

To test if DruE exhibits nuclease activity, we monitored the stabilities of 5-FAM-labeled ssDNA and Alexa Fluor 488-labeled 3′ overhang DNA in the absence or presence of different DruE concentrations. The 3′ overhang DNA substrate was assembled by annealing a 12-nt Alexa Fluor 488-labeled strand and an unlabeled 31-nt strand in a temperature gradient from 95°C to 45°C. 5 pmol of the respective DNA were mixed with 0, 5 or 25 pmol DruE in 40 mM Tris-HCl, pH 7.5, 150 mM NaCl, 0.5 mM MgCl_2_, 4 mM ATP and incubated for 2 h at 37°C. The reactions were terminated using proteinase K. Samples were analyzed by 20% native PAGE. The gels were imaged using a ChemiDoc™ MP Imaging System (BioRad).

### Interaction studies by analytical size exclusion chromatography

Direct DruE-DruH or DruE-DruH-DNA interactions were assessed by analytical SEC on a Superose 6 Increase (3.2/300) column (Cytiva) at 4°C in 10 mM Tris-HCl, pH 8.0, 100 mM NaCl, 5 mM MgCl_2_. DruE and DruH were analyzed separately or after pre-incubation on ice. Additional runs were performed containing a twofold molar excess of forked DNA and/or a fivefold molar excess of various ATP-analogues. Forked DNA was assembled from two single-stranded oligonucleotides with a 13-bp complementary region (**Table S3**).

### Analytical size exclusion chromatography with multi-angle light scattering detection

1 mg/ml DruE or DruE^ΔDD^ were analyzed on a Superose 6 Increase (10/300) column run in 10 mM Tris-HCl, pH 8.0, 100 mM NaCl, 2 mM MgCl_2_ at room temperature. Light scattering was monitored by using a light scattering diode array and a differential refractive index detector (Wyatt Technology). The data were analyzed using ASTRA software (version 6; Wyatt Technology).

### Validation of the DruE-mNeonGreen fusion

The EHEC strains EDL933 Δ*druE* and Δ*druE* pWKS30-*druE*-mNeonGreen were cultivated in LB with 0.2 mM IPTG and harvested after overnight growth by centrifugation, washed in TE-buffer (pH 8.0), disrupted using a RiboLyser (Hybaid), and the protein extracts were cleared from cell debris by centrifugation. The protein extracts were separated using 12% SDS-PAGE, stained with Coomassie blue, and the DruE-mNeonGreen and LacZ bands of the protein extract from Δ*druE* pWKS30-*druE*-mNeonGreen were cut and in-gel digested by trypsin as described ^65^.

Peptides were reconstituted in 10 µl of 0.05% trifluoroacetic acid (TFA) containing 4% acetonitrile, and 2 µl were injected for analysis using an Ultimate 3000 reverse-phase capillary nano-liquid chromatography system coupled to a Q Exactive HF mass spectrometer (Thermo Fisher Scientific). Samples were first loaded onto a trap column (Acclaim PepMap 100 C18, 3 µm, 100 Å, 75 µm i.d. × 2 cm; Thermo Fisher Scientific). After switching the trap column inline, chromatographic separation was performed on an analytical column (Acclaim PepMap 100 C18, 2 µm, 100 Å, 75 µm i.d. × 50 cm; Thermo Fisher Scientific) at a flow rate of 300 nl/min and a column temperature of 55°C. Mobile phase A consisted of 0.1 % formic acid in water, mobile phase B consisted of 0.1% formic acid in 80 % acetonitrile/20 % water. The column was pre-equilibrated with 5% mobile phase B and peptides were separated using a linear gradient from 5% to 44% mobile phase B over 35 min. The mass spectrometer was operated in data-dependent acquisition mode. Survey MS scans were acquired in the range of m/z 350-1650 at a resolution of 60,000, followed by MS/MS scans of the 15 most intense precursor ions at a resolution of 15,000. Dynamic exclusion was set to 20 s. Automatic gain control (AGC) targets were 3 × 10^6^ for MS scans and 1 × 10^5^ for MS/MS scans, with maximum injection times of 20 ms for MS and 25 ms for MS/MS. The isolation window was set to 1.4 m/z and higher-energy collisional dissociation (HCD) was performed with a normalized collision energy of 27. Charge states that were unassigned, singly charged, or higher than 6+ were excluded from fragmentation.

Raw MS and MS/MS data were processed using the Mascot software suite (Mascot Server v3.1, Mascot Daemon v3.0; Matrix Science). Spectra were searched against the proteome of EHEC strain EDL933 downloaded from UniProt (5,520 entries; Proteome ID: UP000028484; January 22, 2026), supplemented with the sequence of the DruE-mNeonGreen protein. Enzyme specificity was set to trypsin, allowing up to two missed cleavages. Precursor and fragment ion mass tolerances were set to 10 ppm and 0.02 Da, respectively. Methionine oxidation and protein N-terminal acetylation were specified as variable modifications. Peptides were considered confidently identified if their Mascot scores exceeded the homology threshold, corresponding to a significance level of p < 0.05 as determined by decoy database searches.

### Plaque assays with *E. coli* MG1655 or ATCC8739 strains

*E. coli* MG1655 with p15a_empty, DruHE or mutated versions of pDruHE, and *E. coli* strain ATCC8739 and deletion mutants were cultivated at 37°C overnight in LB medium. Overnight cultures were diluted 1:100 with LB medium and grown to OD_600_ = 0.5. 2 ml of the cultures were centrifuged at 5,000 x g at 4°C, resuspended in 200 µl of fresh LB medium and stored on ice. 100 µl of the resuspended culture were mixed with soft LB agar (0.5 %) without or with chloramphenicol selection. Selection with ampicillin and chloramphenicol was performed for the complementation assay. Soft LB agar mixed with the bacterial cultures was poured on medium plates with or without antibiotic selection and dried in a clean bench. Phage lysates were diluted 1:10 and spotted in dilutions between 10^-1^ and 10^-8^. The plates were dried and incubated at 37°C overnight. Plaques were counted the next day for plates with dilutions that yielded separated plaques.

Fold protection (FP) and EOP were calculated as log_10_(FP) = a/b and log_10_(EOP) = b/a, respectively, in which a is the number of PFU of *E. coli* MG1655 WT and b is the number of PFU of *E. coli* MG1655 with a vector. The mEOP was calculated as log_10_(mEOP) = a/b - c/d, in which a is the number of PFU of *E. coli* ATCC 8739 Δ*serA/*Δ*rcdB/*Δ*hns/*Δ*stpA*, b is the number of PFU of *E. coli* ATCC8739, c is the number of PFU of *E. coli* ATCC8739 Δ*druHE* Δ*serA/*Δ*rcdB/*Δ*hns/*Δ*stpA*, and d is the number of PFU of *E. coli* ATCC8739 Δ*druHE*.

### Plaque assays with EDL933 strains

The pWKS30-*druE* variant and mNeonGreen fusion plasmids were electroporated into the Δ*druE* mutant, generating strains EDL933 Δ*stx1/2* Δ*druE* pWKS30-*druE* variants and pWKS30-*druE-*mNeonGreen, which were screened for ampicillin resistance. EOP assays were performed to determine the functionality of the pWKS30-encoded DruE variants in the defense against phage Bas26 of the BASEL collection^33^. For EOP assays, square Petri dishes (12 cm × 12 cm) were used. 9 ml of top agar (0.5% agar, 20 mM MgSO_4_, 5 mM CaCl_2_) were inoculated with 200 μl of overnight culture of the EDL933 WT strain, the Δ*druE* mutant, or WT *druE* and *druE* variant-complemented strains to overlay the solid LB agar plates. Afterwards, 5 μl of serial dilutions of phage Bas26 lysate stock were spotted onto the top agar plates and incubated at 37°C for 24 h to determine plaque formation. The EOP of Bas26 was quantified by calculating the ratio of plaques of the EDL933 WT and the complemented strains expressing pWKS30-*druE* or pWKS30-*druE* variants versus the Δ*druE* mutant strain. The EOP assays were performed in 3-4 biological replicates.

### Liquid culture assays with *E. coli* MG1655 strains

*E. coli* MG1655 with p15a_empty or pDruHE were cultivated at 37°C overnight in LB medium with antibiotic selection. The overnight cultures were diluted 1:100 in LB medium and grown to OD_600_ = 0.3. 180 µl of the cultures were mixed in Flat-Bottom Nuclon 96-well plates with 20 µl of phage lysate at concentrations to reach MOIs of 10, 5, 1, 0.5, 0.1, or 0.05. The 96-well plates were incubated at 37°C for 16 h with orbital rotation in a Synergy Multimode Reader (BioTek) and OD_600_ values were recorded every 3 min.

### Phage adsorption assay

*E. coli* MG1655 with p15a_empty or pDruHE were cultivated at 37°C overnight in LB medium. The overnight cultures were diluted 1:100 in LB medium and grown to OD_600_ = 0.3. 180 µl of the cultures were mixed in Flat-Bottom Nuclon 96-well plates with 20 µl of phage lysate to reach an MOI of 1. The 96-well plates were incubated at 37°C with orbital rotation in a Synergy Multimode Reader. 100 µl of the cultures were harvested every 10 min during the first h and at 2 and 3 h post-infection. Samples were immediately centrifuged at 5,000 x g for 5 min and plated on soft LB agar with the propagation strain *E. coli* BW25113 as described above.

### Mitomycin C toxicity assay

*E. coli* MG1655 with the empty vector control (p15a_empty) or pDruHE, or pDruHE^E525A^ were cultivated at 37°C overnight in LB medium. The overnight cultures were diluted 1:100 in LB medium and grown to OD_600_ = 0.3. Cultures were exposed to 5 µg/ml mitomycin C and incubated for an additional 30 min. The cultures were serially diluted and spotted on LB agar plates containing 5µg/ml mitomycin C. Plates were dried and incubated at 37°C overnight. Colonies were counted the next day. Mitomycin C toxicity was calculated as log_10_(-Δ) = a/b - c/d, in which a = CFU of the strain with the defense system with mitomycin C; b = CFU of the strain with the defense system without mitomycin C; c = CFU of the strain without the defense with mitomycin C; d = CFU of the strain without the defense without mitomycin.

### Mass spectrometry analysis

*E. coli* ATCC8739 Δ*druHE* pDruHE-3F and ATCC8739 Δ*druHE* pDruHE were cultivated at 37°C overnight in LB medium. The overnight cultures were diluted 1:100 in 400 ml of LB medium and grown to OD_600_ = 0.3. Phage Bas23 lysate was added at a MOI of 10 or not, and cultures were incubated for 5 min without rotation at 37°C, then for an additional 5 min with rotation. The cultures were centrifuged at 4,000 x g for 10 min at 4°C, resuspended in 10 ml phosphate-buffered saline (PBS), washed a second time, resuspended in 1 ml PBS, washed a third time, and resuspended in 500 µl PBS. The samples were incubated at 30°C with 500 µg/ml of lysozyme and 30 mM of NaCl for 1 h, centrifuged at 8,000 x g for 10 min at 4°C, and the supernatant was recovered. DruE-3F was immunoprecipitated with Pierce™ Anti-DYKDDDDK Magnetic Agarose (Thermo Fisher Scientific) overnight at 4°C, and the beads were washed using the protocol and the washing buffer from the Dynabeads™ Protein A Immunoprecipitation Kit (Thermo Fisher Scientific). DruE-3F and bound proteins were eluted with Pierce™ 3x DYKDDDDK Peptide (Thermo Fisher Scientific) at 0.5 mg/ml in PBS overnight at 4°C.

Mass spectrometry and data analysis were carried out by the proteomics core facility of the European Molecular Biology Laboratory in Heidelberg using TMT-based quantitative proteomics. Protein samples were analyzed *via* the SP3 protocol^66^ on a KingFisher Apex™ platform (Thermo Fisher Scientific). Samples were digested with trypsin in a 1:20 ratio (protease:protein) in 50 mM 4-(2-hydroxyethylpiperazine) For the proteomics data, 4-(2-hydroxyethyl)-1-piperazineethanesulfonic acid (HEPES) supplemented with 5 mM Tris(2-carboxyethyl)phosphine hydrochloride (TCEP) and 20 mM 2-chloroacetamide (CAA) for 5 h at 37 °C. Up to 10 µg of peptides were labeled using TMTpro™ 16-plex reagent (Therno Fisher Scientific) as previously described^67^. Briefly, 0.5 mg of TMT reagent was dissolved in 45 µl of 100 % acetonitrile. 4 µl of the solution were added to each peptide sample, followed by incubation for 1 h at room temperature. The labeling reaction was quenched by adding 4 µl of a 5 % aqueous hydroxylamine solution and incubating for an additional 15 min at room temperature. Labeled samples were combined for multiplexing, desalted using an Oasis® HLB µElution Plate (Waters) according to the manufacturer’s instructions, and dried by vacuum centrifugation.

Offline high-pH reversed-phase fractionation^68^ was carried out using a 1200 Infinity high-performance liquid chromatography (HPLC) system (Agilent), equipped with a Gemini C18 analytical column (3 μm particle size, 110 Å pore size, dimensions 100 x 1.0 mm; Phenomenex) and a Gemini C18 SecurityGuard pre-column cartridge (4 x 2.0 mm; Phenomenex). The mobile phases consisted of 20 mM ammonium formate, pH 10.0 (buffer A), and 100% acetonitrile (buffer B). The peptides were separated at a flow rate of 0.1 ml/min using the following linear gradient: 100% buffer A for 2 min, ramping to 35 % buffer B over 59 min, increasing rapidly to 85 % buffer B within 1 min, and holding at 85 % buffer B for an additional 15 min. Subsequently, the column was returned to 100% buffer A and re-equilibrated for 13 min. During the LC separation, 48 fractions were collected and pooled in groups of 6. Pooled fractions were dried using vacuum centrifugation.

An UltiMate 3000 RSLCnano LC system (Thermo Fisher Scientific) equipped with a trapping cartridge (µ-Precolumn C18 PepMap™ 100, 300 µm i.d. × 5 mm, 5 µm particle size, 100 Å pore size; Thermo Fisher Scientific) and an analytical column (nanoEase™ M/Z HSS T3, 75 µm i.d. × 250 mm, 1.8 µm particle size, 100 Å pore size; Waters). Samples were trapped at a constant flow rate of 30 µl/min using 0.05% trifluoroacetic acid (TFA) in water for 6 min. After switching in-line with the analytical column, which was pre-equilibrated with solvent A (3% dimethyl sulfoxide (DMSO), 0.1% formic acid in water), the peptides were eluted at a constant flow rate of 0.3 µl/min using a gradient of increasing solvent B concentration (3% DMSO, 0.1% formic acid in acetonitrile) The gradient was as follows: 2% to 8% solvent B in 6 min, 8% to 25% solvent B in 69 min, 25% to 40% solvent B in 5 min, 40% to 85% solvent B in 0.1 min, maintenance at 85% solvent B for 3.9 min, and a re-equilibration to 2% solvent B for 6 min.

Peptides were introduced into an Orbitrap Fusion™ Lumos™ Tribrid™ mass spectrometer (Thermo Fisher Scientific) *via* a Pico-Tip emitter (360 µm OD × 20 µm ID; 10 µm tip; CoAnn Technologies) using an applied spray voltage of 2.2LJkV. The capillary temperature was maintained at 275°C. Full MS scans were acquired in profile mode over an m/z range of 375 - 1,500, with a resolution of 120,000 at m/z 200 in the Orbitrap. The maximum injection time was set to 50LJms, and the AGC target limit was set to “standard”. The instrument was operated in data-dependent acquisition (DDA) mode, with MS/MS scans acquired in the Orbitrap at a resolution of 30,000. The maximum injection time was set to 94LJms, with an AGC target of 200%. Fragmentation was performed using HCD with a normalized collision energy of 34%, and MS2 spectra were acquired in profile mode. The quadrupole isolation window was set to 0.7 m/z, and dynamic exclusion was enabled with a duration of 60 s. Only precursor ions with charge states 2–7 were selected for fragmentation.

Raw files were converted to mzML format using MSConvert (ProteoWizard), using peak picking, 64-bit encoding, and zlib compression, and filtering for the 1000 most intense peaks. Files were then searched using MSFragger in FragPipe (version 23.0) against FASTA databases Bas23_phageDB_v2 and P4266_E.coli_ATCC8739_v2 containing common contaminants and reversed sequences. The following modifications were included into the search parameters: Carbamidomethylation (C, 57.0215) and TMTpro (K, 304.2072) as fixed modifications; oxidation (M, 15.9949), acetylation (protein N-terminus, 42.0106), and TMTpro (peptide N-terminus, 304.2072) as variable modifications. For the full scan (MS1) a mass error tolerance of 20 ppm and for MS/MS (MS2) spectra of 20 ppm was set. For protein digestion, “trypsin” was used as protease with an allowance of maximum 2 missed cleavages requiring a minimum peptide length of 7 residues. The false discovery rate (FDR) on peptide and protein level was set to 0.01.

For the proteomics data analysis, the raw output files of FragPipe (protein.tsv files files) were processed using the R programming environment. Initial data processing included filtering out contaminants and reverse proteins. Only proteins quantified with at least 2 razor peptides (with Razor.Peptides >= 2) were considered for further analysis. 1,155 proteins passed the quality control filters. To correct for technical variability, batch effects were removed using the “removeBatchEffect” function of the limma package^69^ on the log_2_-transformed raw TMT reporter ion intensities (“channel” columns). Subsequently, normalization was performed using the “normalizeVSN” function of the limma package (VSN - variance stabilization normalization)^70^. Differential expression analysis was performed using the moderated t-test provided by the limma package^69^. The model accounted for replicate information by including it as a factor in the design matrix passed to the “lmFit” function. Proteins were annotated as hits if they had an FDR below 0.05 and an absolute fold change greater than 1.5. Proteins were considered candidates if they had an FDR below 0.2 and an absolute fold change greater than 1.5. Clustering with all enriched hit and enriched candidate proteins based on the median protein abundances normalized by median of control condition was conducted to identify groups of protein with similar patterns across conditions. The “kmeans” method was employed, using Euclidean distance as the distance metric and “ward.D2” linkage for hierarchical clustering. The optimal number of clusters (6) was determined using the Elbow method, which identifies the point where the within-group sum of squares stabilizes.

### Total internal reflection (TIRF) microscopy of DruE-mNeonGreen

Overnight cultures of the EHEC strain EDL933 Δ*druE* pWKS30-*druE*-mNeonGreen, encoding a DruE-mNeonGreen translational fusion, were grown in LB medium without glucose at 37 °C with shaking (180 rpm) and supplemented with the appropriate antibiotics. The next day, cultures were diluted 1:100 into 10 ml of fresh LB supplemented with antibiotics, 20 mM MgSO_4_, 5 mM CaCl_2_, and 0.2 mM IPTG, and cultivated at 37°C until mid-exponential phase (OD_600_ = 0.4-0.6). For the exposure of cells to the phages, cultures were normalized to an OD_600_ of 0.2 by dilution into supplemented LB as stated above. Normalized cultures were either left untreated (no-phage control) or infected with bacteriophage Bas67 at MOI = 100. Infections were carried out in 2 ml centrifuge tubes under shaking conditions (< 650 rpm) in a ThermoMixer at 37°C. Samples were collected at 1, 2, and 3 h post infection. For TIRF microscopy, 1 µl of cells and phage mix was spotted onto an agarose pad (1.2 % in Milli-Q water of UltraPure agarose; Invitrogen) and directly imaged. TIRF microscopy was performed using an Eclipse Ti2 inverted microscope (Nikon) equipped with an ILAS 2 TIRF module (Gataca Systems) and a TIRF × 100/1.49 NA oil objective. The samples were excited with a 50-ms exposure using a 561 nm laser at 80%, and emission was recovered through a quad TIRF filter cube (emission: 525/50 nm).

### Northern blotting and motility assays of EDL933 Δ*druE* mutant and complemented strain

For *fliC* transcriptional analysis, the EDL933 Δ*druE* mutant and the *druE*+ complemented strains were cultivated in LB medium and harvested at an OD_600_ of 0.8 for RNA isolation, using the acid phenol extraction method as previously described^71^. Northern blot hybridizations were conducted with digoxigenin-labeled *fliC*-specific antisense RNA probes, synthesized by *in vitro* transcription using T7 RNA polymerase as described previously^71^. Swimming assays were performed to analyze motility of the EDL933 Δ*druE* mutant and *druE*+ complemented strain. Overnight cultures of the *E. coli* strains (3 µl of OD_600_ = 1) were spotted on LB plates containing 0.3% soft agar. The diameters of swimming zones were quantified after incubation at 37 °C for 12 h.

### CryoEM sample preparation

Forked DNA was assembled using two single-stranded oligonucleotides with a 13 bp complementary region (**Table S3**). Both strands were mixed in a 1:1 ratio, heated to 90 °C for 2 min, and slowly cooled to room temperature. Purified DruE was mixed with a 1.5 molar excess of the DNA substrate, 2 mM (final) ATPүS, and 2 mM (final) MgCl_2_. The mixture was incubated on ice for 30 min. n-octyl-β-D-glucoside (0.15 % [v/v] final concentration) was added to the mixture immediately before vitrification. 3.8 μl of the final sample were applied to glow-discharged Quantifoil R1.2/1.3 holey carbon grids and plunged into liquid ethane using a Vitrobot Mark IV (Thermo Fisher Scientific) set at 10°C and 100% humidity.

### CryoEM data acquisition and analysis

CryoEM data were acquired on an FEI Titan Krios G3i TEM (Thermo Fisher Scientific) operated at 300 kV and equipped with a Falcon 3EC direct electron detector (Thermo Fisher Scientific). Movies were recorded for 40.57 s accumulating a total electron flux of 40 e^-^/Å^2^ in counting mode distributed over 33 fractions at a nominal magnification of 96,000x, yielding a calibrated pixel size of 0.832 Å/px for DruE. For DruH a magnification of 120,000x was selected, yielding a calibrated pixel size of 0.657 Å/px.

Automated data acquisition was conducted using EPU software (Thermo Fisher Scientific). All image analysis steps were carried out using cryoSPARC^72^. Movie alignment was done with patch motion correction, CTF estimation was conducted with Patch CTF. Class averages of manually selected particle images were used to generate an initial template for reference-based particle picking from 2,742 micrographs. 660,861 particle images were extracted with a box size of 384 px and Fourier-cropped to 96 px for initial analysis. Reference-free 2D classification was used to select 345,647 particle images for further analysis. *Ab initio* reconstruction was conducted to generate an initial 3D reference for heterogeneous 3D refinement, after which 469,099 particles were re-extracted with a box size of 384 px Fourier-cropped to 288 px. After multiple iterations of heterogeneous refinement, one monomer structure and five dimer structures were selected for non-uniform (NU) refinement, yielding final reconstructions at global resolutions between 2.9 Å and 4.2 Å. Local resolutions range from 2.5-3.0 Å in core regions to 8.5-11.0 Å in some peripheral elements (**Figures S4 and S5**; **Table S1**).

CryoEM data for the DruH sample were analyzed accordingly with minor modifications. 67,184 particle images from 1,863 micrographs were used for multi-particle 3D refinement of the DruH reconstruction. Before final NU refinement, selected particles were subjected to local motion correction and extraction with a box size of 384 px (without Fourier-cropping), yielding a reconstruction at 3.3 Å global resolution, with local resolution ranging from 2.5 to 8.5 Å (**Figures S6 and S7**; **Table S1**)

### Model building, refinement, and analysis

Initially the model of monomeric DruE was manually built into the cryoEM reconstruction. Model building was guided by the structure of the MfrA helicase (PDB ID: 6ZNS)^27^. The initial model was adjusted by rigid body fitting and segmental real-space refinement using Coot (version 0.9.6)^73,74^. For model building of the dimeric structures, individual domains of monomeric DruE were placed into the cryoEM reconstructions. While we could unequivocally trace the DNA strands in the cryoEM reconstructions of the DruE-ATPγS-DNA complexes, the resolutions obtained did not allow assignment of the nature of all DNA nucleotides. In the monomeric complex, we placed the DNA substrate based on well-defined density for the 3′ end of the tracking strand, which we assigned as residue Thy30 of the tracking strand. This assignment left eleven bp in the duplex region. In the state IV protomer of DurE dimer 1 (**Figure 4B**), the DNA was placed based on the well-defined density of the duplex region. This assignment led to a 12-bp duplex and a register shift along the 3′ overhang compared to the monomeric complex. Models were refined by iterative rounds of real space refinement in PHENIX (version 1.21_5419)^75,76^ and manual adjustment in Coot. The final structural models were evaluated with MolProbity (version 4.5.2) ^77^. Structure figures were prepared with PyMOL (version 3.1.2; Schrödinger) and ChimeraX ^78,79^. DruE regions 524-533, 1356-1365, and 2031-2037 lack well-defined density in most or all reconstructions, density for residues 1897-2104 is very weak or missing in several reconstructions, and density for region 681-757 is lacking from the reconstruction for the monomeric DruE complex.

For DruH a model predicted by AlphaFold^80^ was docked into the cryoEM reconstruction using PHENIX, manually adjusted, and refined as for the DruE complexes. Residues 7-1117 of DruH could be unambiguously fitted to the cryoEM reconstruction.

