## Supplementary Information for "A compact Druantia defense clears phage infections *via* single-stranded DNA recognition and directional duplex unwinding"

#### SUPPLEMENTAL FIGURES

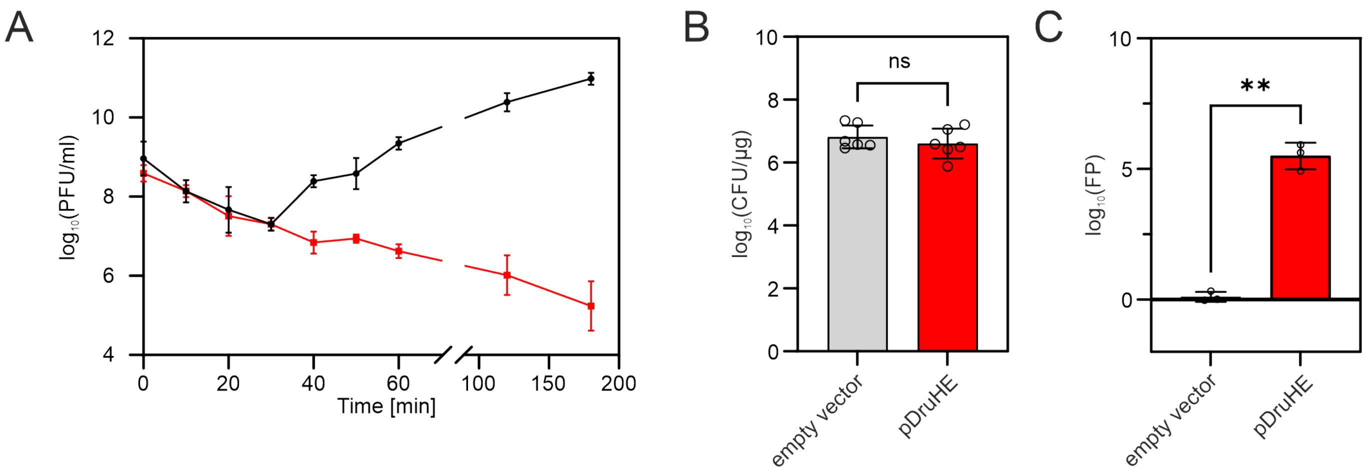

**Figure S1. Adsorption assay and methylation-free *E. coli* strain. Related to Figure 1.**

(A) Adsorption of Bas29 on *E. coli* MG1655 with an empty vector (black line) or with pDruHE (red line), measured as PFU/ml with plaque assay of the supernatant. Filled shapes and error bars represent the log-transformed mean ± SD for three biological replicates.

(B) Transformation efficiency of the *E. coli* EC135 strain with the empty vector or pDruHE. Bars and error bars represent the mean ± SD of log-transformed values for six biological replicates. Open circles represent individual experiments.

(C) Sensitivity of *E. coli* EC135 producing Druantia III-A to Bas29, compared to *E. coli* EC135 transformed with an empty vector. Data were acquired *via* a plaque assay and sensitivity is represented as fold protection (FP). Bars and error bars represent the mean ± SD from three biological replicates. Open circles represent individual experiments.

Welch's t-test was used to assess significance. **, $p \leq 0.01$; n.s., not significant.

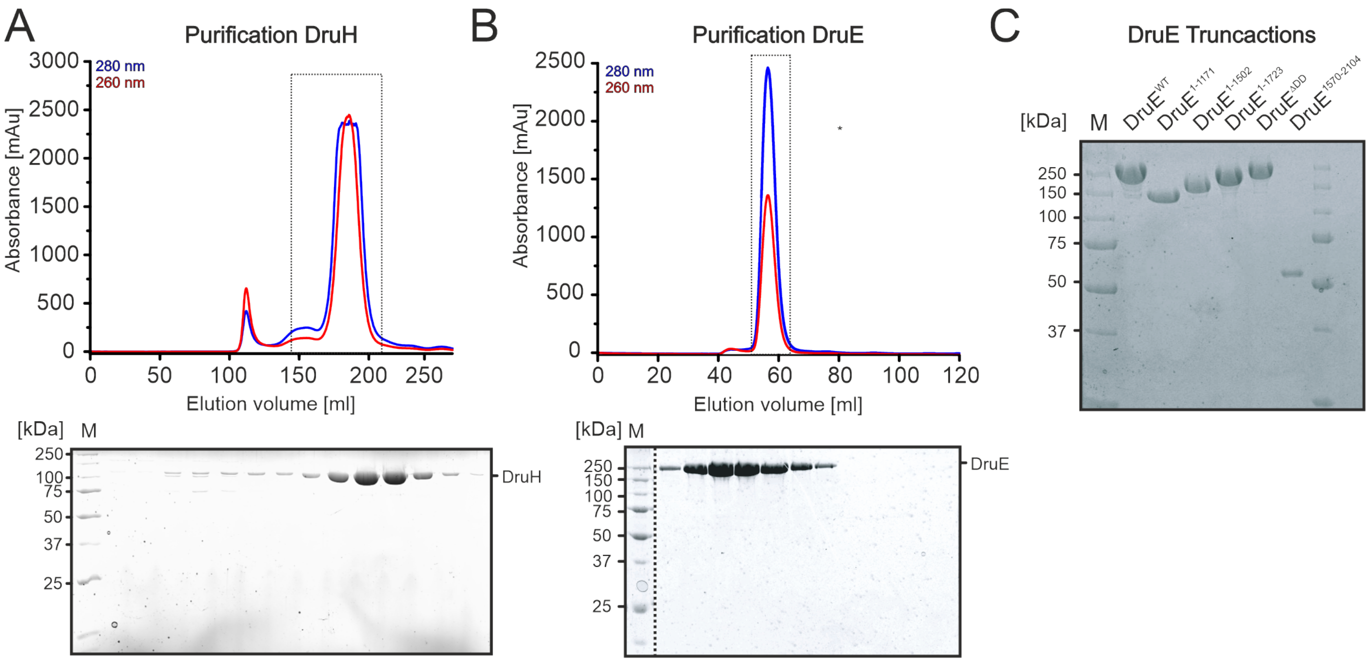

**Figure S2. Overview of protein purification. Related to Figures 2-6.**

(A) UV traces (top panel) and SDS-PAGE analysis (bottom panel) of the peak fraction of the final SEC run of a DruH purification. Boxed region in top panel, fractions analyzed by SDS-PAGE. M, marker.

(B) UV traces (top panel) and SDS-PAGE analysis (bottom panel) of the peak fraction of the final SEC run of a DruE purification. Boxed region in top panel, fractions analyzed by SDS-PAGE. The SDS gel was spliced to remove irrelevant lanes. Dashed line, splice position. M, marker.

(C) SDS-PAGE analysis of purified DruE variants (DruE$^{1\text{-}1171}$, DruE$^{1\text{-}1502}$, DruE$^{1\text{-}1723}$, DruE$^{1570\text{-}2104}$, and DruE$^{\Delta DD}$). M, marker.

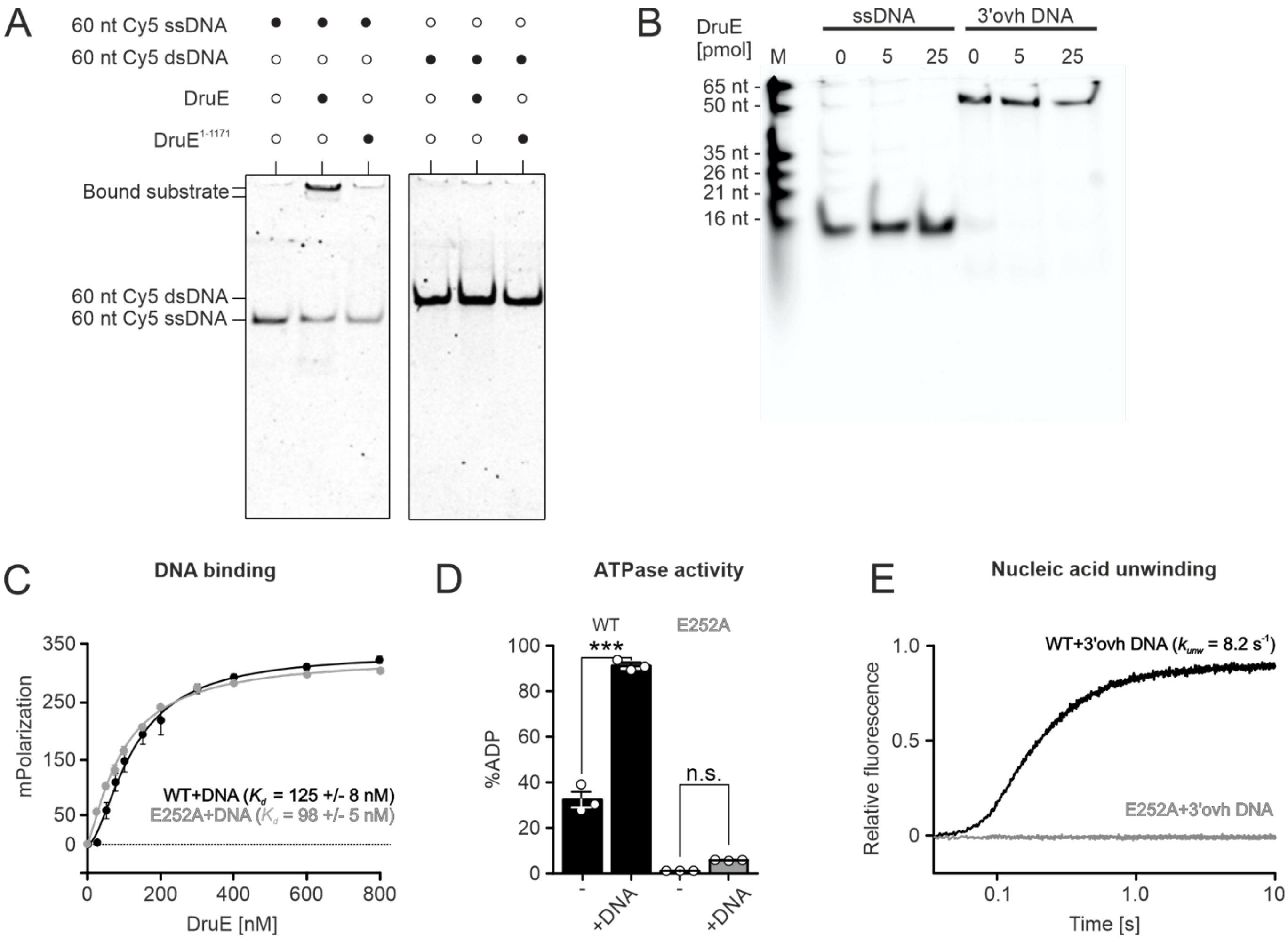

**Figure S3.** ***In vitro*** **characterization of DruE. Related to Figure 2.**

(A) Electrophoretic mobility shift assay monitoring binding of the indicated DruE variants to ssDNA and dsDNA substrates. 60 nt Cy5 dsDNA, 5' Cy5-labeled 60-nt double-stranded DNA; 60 nt Cy5 ssDNA, 5' Cy5-labeled 60-nt single-stranded DNA.

(B) Native PAGE analysis monitoring stabilities of DNA substrates used in binding and unwinding experiments in the presence of increasing concentrations of $DruE^{WT}$.

(C) FA assay monitoring binding of the indicated DruE variants to a 15-nt ssDNA. Data represent means ± SEM of three independent experiments using the same biochemical samples.

(D) Intrinsic and DNA-stimulated ATPase activities of the indicated DruE variants derived from thin-layer chromatographic analyses. Data represent means ± SEM of three independent experiments using the same biochemical samples. Open circles indicate individual experiments. Significance was assessed *via* unpaired Student's-t-tests. *, $p \leq 0.05$; **, $p \leq 0.01$; ns, not significant.

(E) Unwinding of 3′ overhang (ovh) DNA by the indicated DruE variants monitored *via* stopped-flow/fluorescence assays. Single representative traces of three independent measurements are shown.

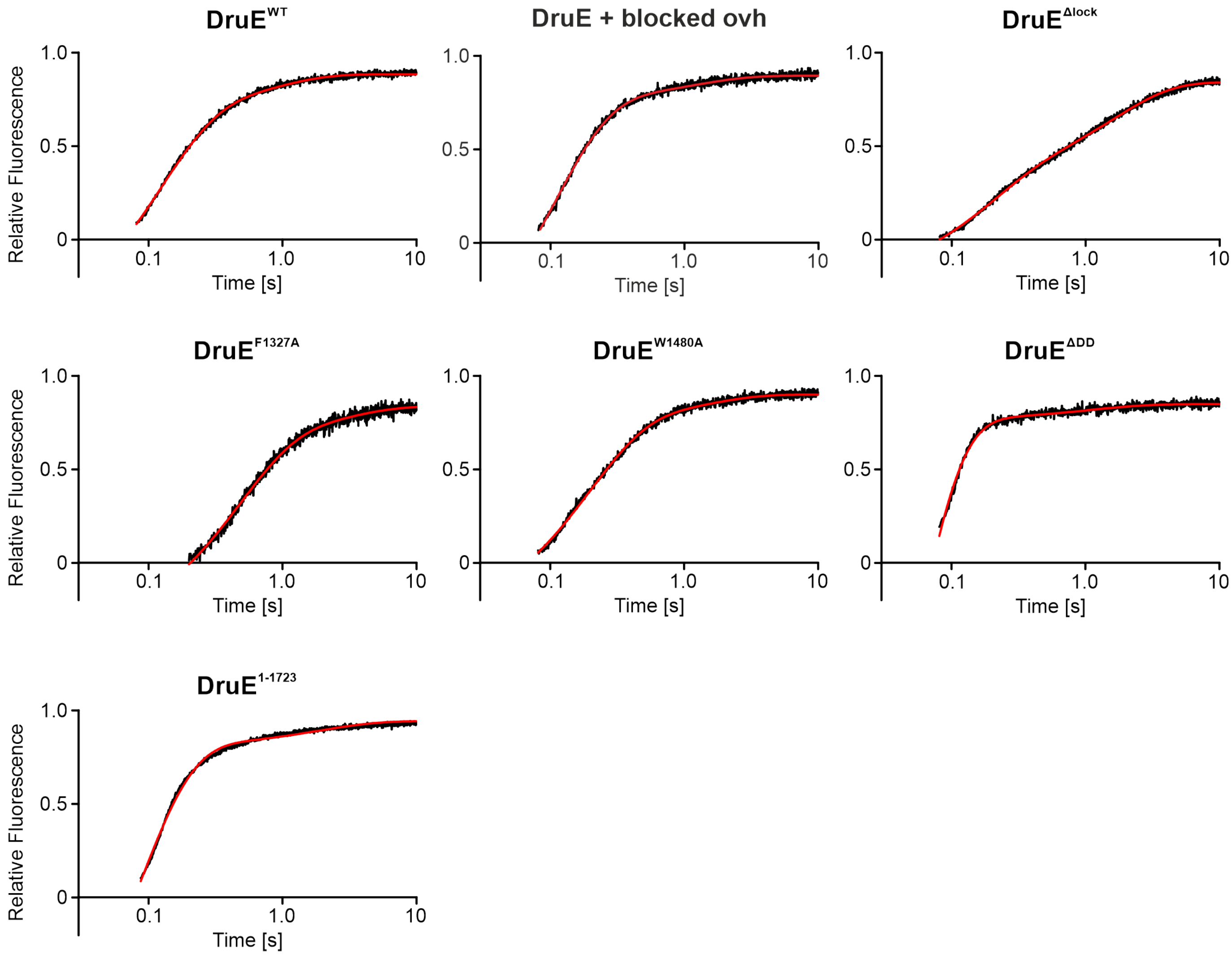

**Figure S4. Fitting of stopped-flow/fluorescence data. Related to Figures 2-6.**

The data were normalized and fitted to a double exponential equation (fraction unwound = $A_{fast}\cdot$(1-exp(-$k_{fast}$t))+$A_{slow}\cdot$(1-exp(-$k_{slow}$t)); $A_{fast/slow}$, unwinding amplitude of the fast/slow phase; $k_{fast/slow}$, unwinding rate constants of the fast/slow phase [$s^{-1}$]; t, time [s]). The first 0.08 s (0.2 s for DruE$^{F1327A}$) of data acquisition were excluded from curve fitting to account for the initial mixing periods. Amplitude-weighted unwinding rate constants were calculated as $k_{unw} = \Sigma(A_i k_i^2)/\Sigma(k_i A_i)$.

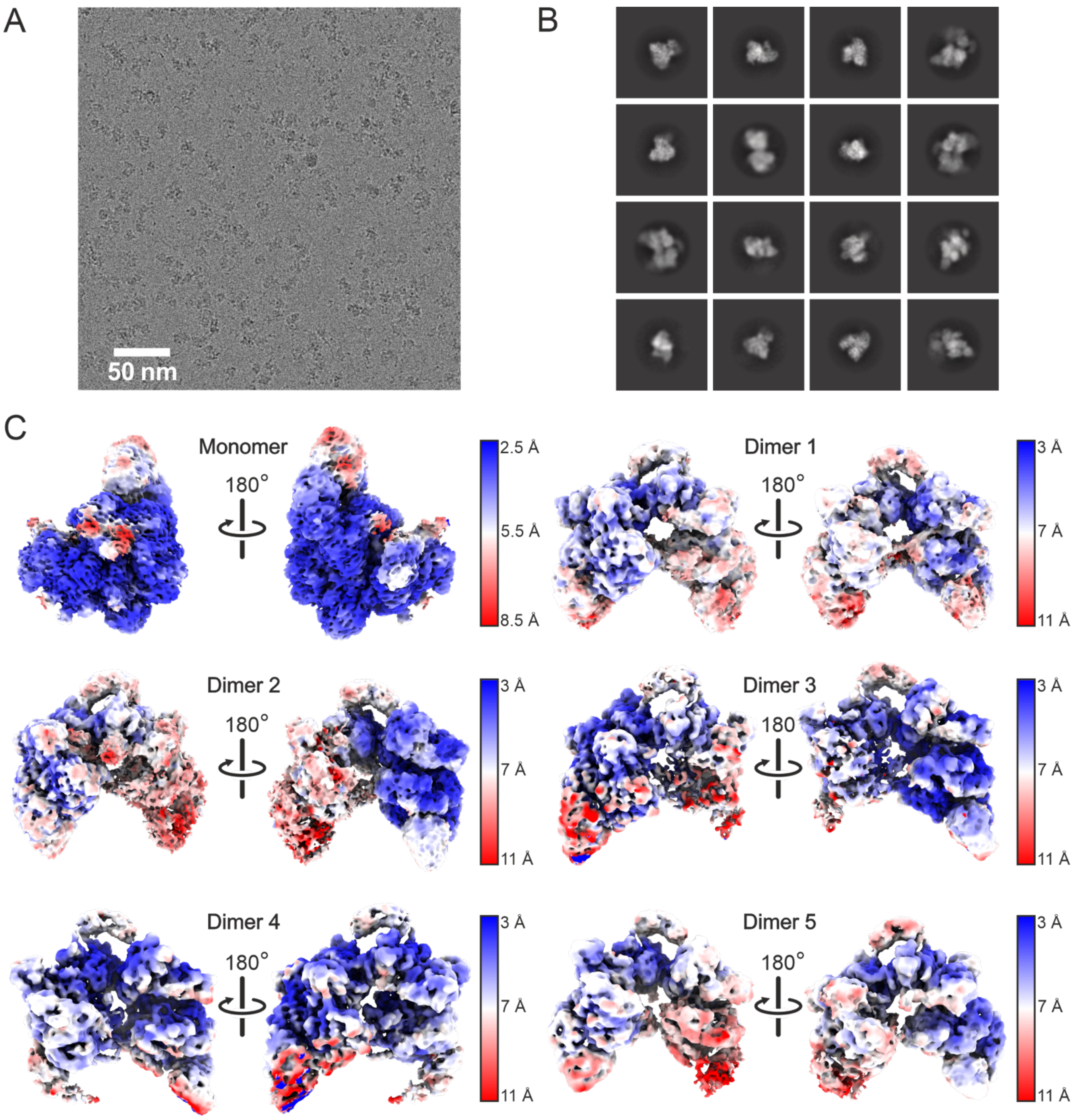

**Figure S5. DruE cryoEM data. Related to Figure 3.**

(A) Representative cryoEM micrograph of DruE. Scale bar, 50 nm.

(B) Selected 2D class averages after reference-free 2D classification of DruE-ATPγS-DNA complexes.

(C) Diametric views of cryoEM reconstructions of the monomeric and the five dimeric DruE-ATPγS-DNA complexes, colored by local resolution as indicated in the legends.

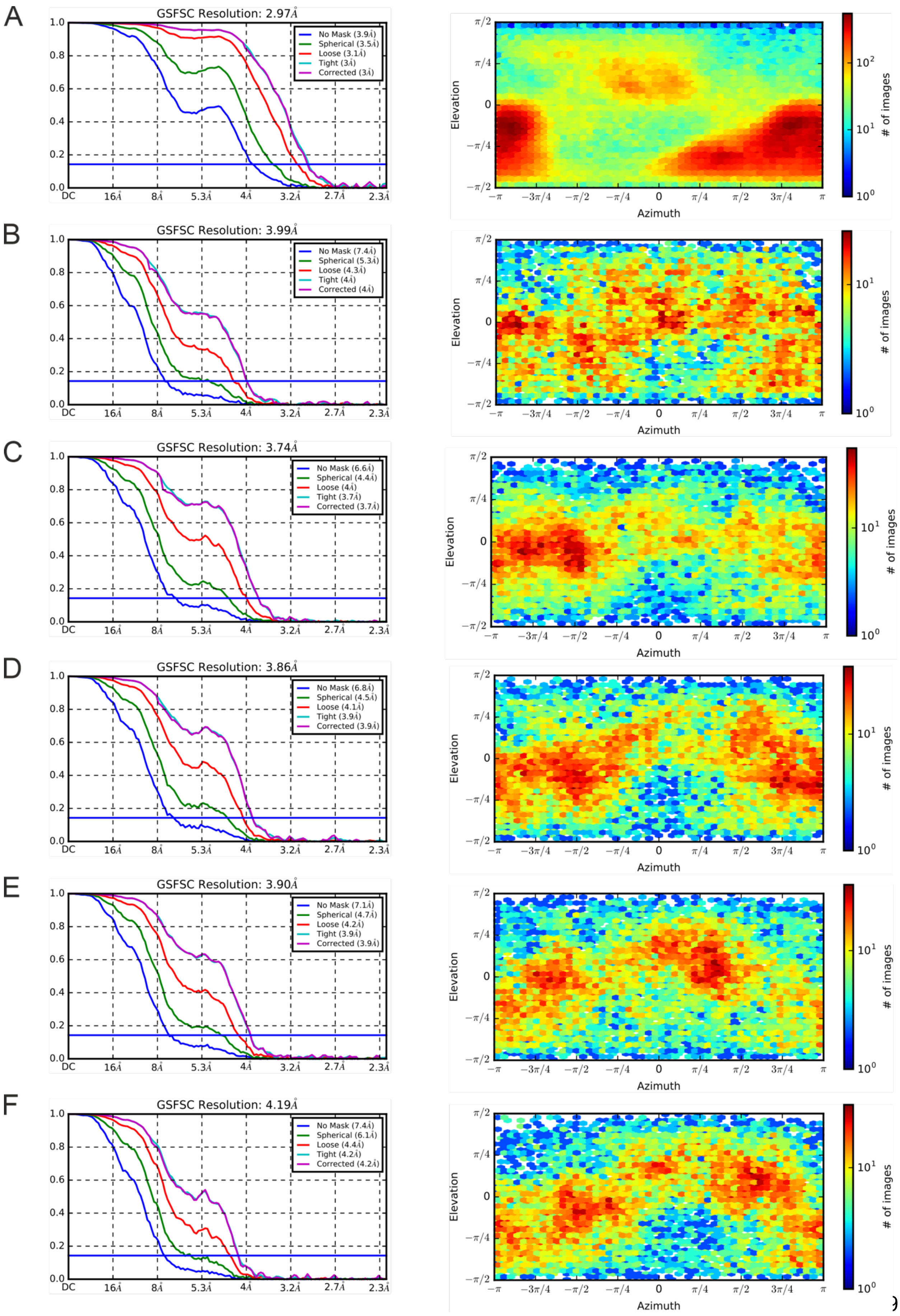

A
GSFSC Resolution: 2.97Å
No Mask (3.9Å)
Spherical (3.5Å)
Loose (3.1Å)
Tight (3Å)
Corrected (3Å)
B
GSFSC Resolution: 3.99Å
No Mask (7.4Å)
Spherical (5.3Å)
Loose (4.3Å)
Tight (4Å)
Corrected (4Å)
C
GSFSC Resolution: 3.74Å
No Mask (6.6Å)
Spherical (4.4Å)
Loose (4Å)
Tight (3.7Å)
Corrected (3.7Å)
D
GSFSC Resolution: 3.86Å
No Mask (6.8Å)
Spherical (4.5Å)
Loose (4.1Å)
Tight (3.9Å)
Corrected (3.9Å)
E
GSFSC Resolution: 3.90Å
No Mask (7.1Å)
Spherical (4.7Å)
Loose (4.2Å)
Tight (3.9Å)
Corrected (3.9Å)
F
GSFSC Resolution: 4.19Å
No Mask (7.4Å)
Spherical (6.1Å)
Loose (4.4Å)
Tight (4.2Å)
Corrected (4.2Å)
DC
16Å
8Å
5.3Å
4Å
3.2Å
2.7Å
2.3Å
Elevation
Azimuth
### of images

**Figure S6. DruE cryoEM data. Related to Figure 3.**

Gold standard Fourier shell correlation analysis and viewing direction distribution of monomeric DruE-DNA-ATPγS (A), DruE-DNA-ATPγS, dimer 1 (B), DruE-DNA-ATPγS, dimer 2 (C), DruE-DNA-ATPγS, dimer 3 (D), DruE-DNA-ATPγS, dimer 4 (E), and DruE-DNA-ATPγS, dimer 5 (F).

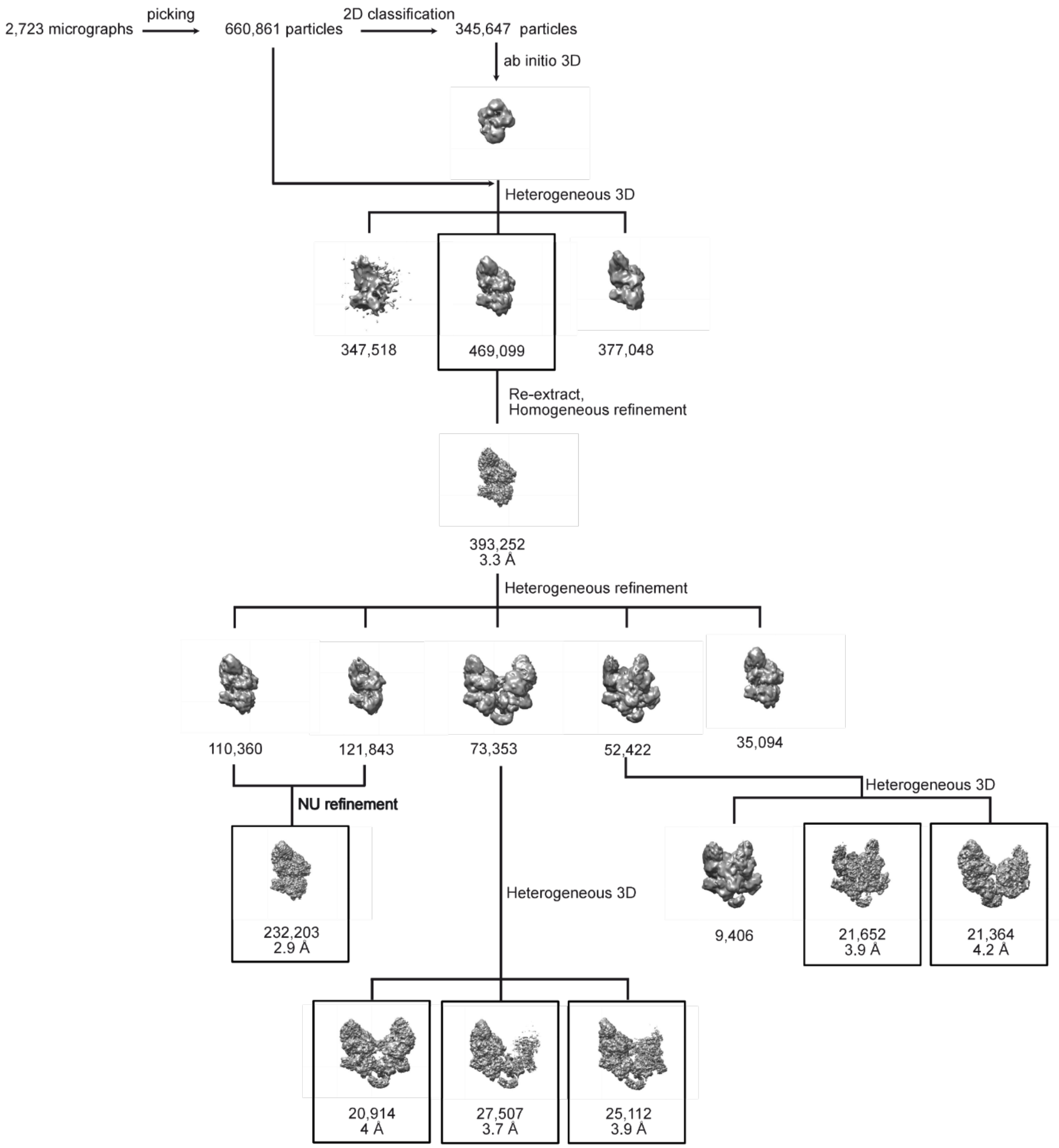

**Figure S7. DruE hierarchical clustering analysis. Related to Figure 3.**

660,861 particle images were picked from 2,723 micrographs, extracted with Fourier cropping, and subjected to refence-free 2D classification, after which a subset of 345,647 particle images was selected for *ab initio* 3D reconstruction. The obtained map was used as a reference for heterogenous 3D refinement of the entire dataset to select 469,099 particle images for re-extraction without binning. Further iterative heterogenous refinement revealed separation into monomeric and dimeric species, final reconstructions were obtained by non-uniform (NU) refinement with global resolutions between 2.9 Å and 4.2 Å as indicated.

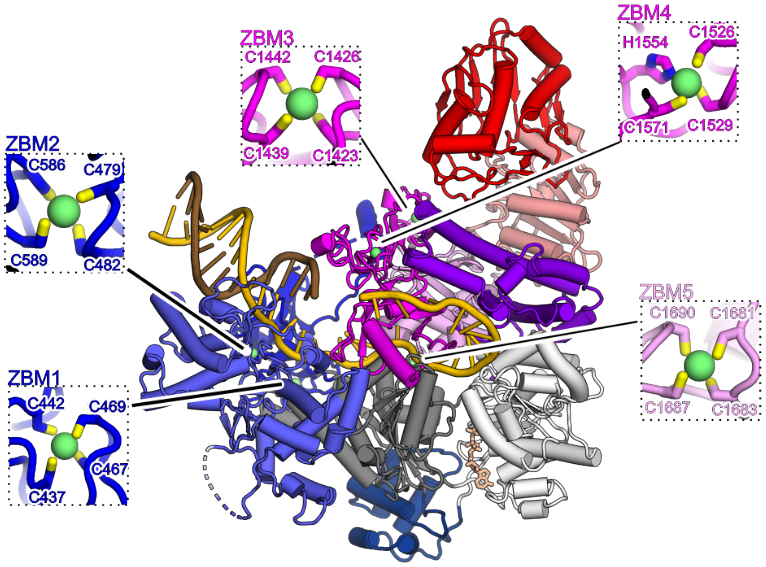

**Figure S8. DruE zinc-binding motifs. Related to Figure 3.**

DruE comprises five ZBMs in the dZBD (ZBM1 and 2), the OBZ (ZBM3 and 4) and MZB (ZBM5) domains. Enlarged views on the ZBMs show the zinc-coordinating residues in stick representation colored by atom type; carbon, as the respective domain; nitrogen, blue; sulfur, yellow.

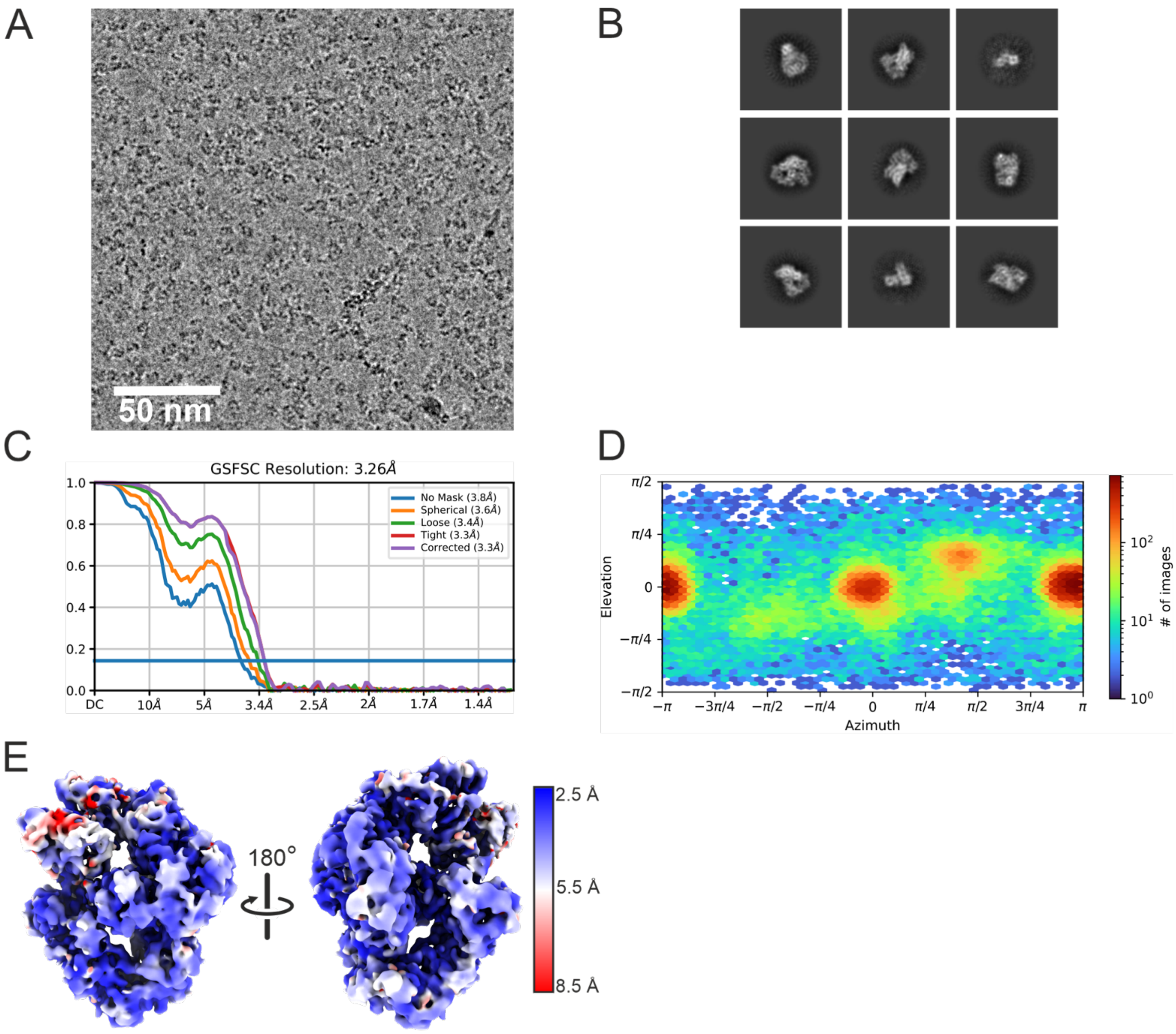

**Figure S9. DruH cryoEM data. Related to Figure 6.**

(A) Representative cryoEM micrograph. Scale bar, 50 nm.

(B) Selected 2D class averages after reference-free 2D classification of DruH.

(C) Gold standard Fourier shell correlation analysis of DruH.

(D) Viewing direction distribution of DruH.

(E) Diametric views of the DruH cryoEM reconstruction, colored by local resolution as indicated in the legend.

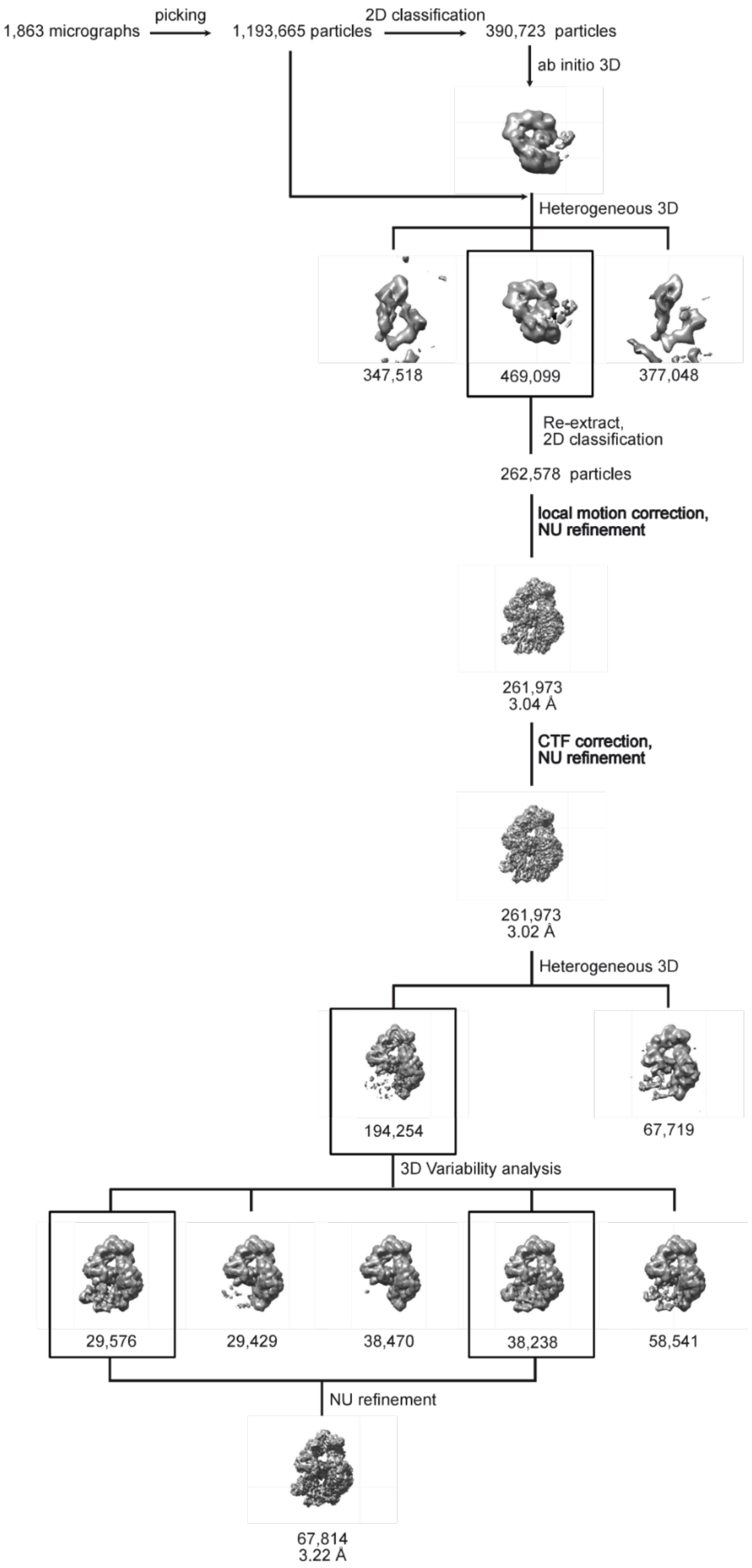
1,863 micrographs
picking
1,193,665 particles
2D classification
390,723 particles
ab initio 3D
Heterogeneous 3D
347,518
469,099
377,048
Re-extract,
2D classification
262,578 particles
local motion correction,
NU refinement
261,973
3.04 Å
CTF correction,
NU refinement
261,973
3.02 Å
Heterogeneous 3D
194,254
67,719
3D Variability analysis
29,576
29,429
38,470
38,238
58,541
NU refinement
67,814
3.22 Å

**Figure S10. DruH hierarchical clustering analysis. Related to Figure 6.**

1,193,665 particle images were picked, extracted with Fourier cropping from 1,863 micrographs, and subjected to reference-free 2D classification. A reference obtained by *ab initio* 3D reconstruction from a subset of 390,723 particle images was used for heterogenous 3D refinement of the entire dataset into three classes. The selected particle images were re-extracted without Fourier cropping and subjected again to 2D classification, after which 262,579 particle images were selected for further processing including local motion correction, CTF refinement, and NU refinement. Fragmented density in the consensus reconstruction was improved by 3D variability analysis using a focus mask on the fragmented region, from which a final subset of 67,814 particle images was selected for NU refinement to a global resolution of 3.22 Å.

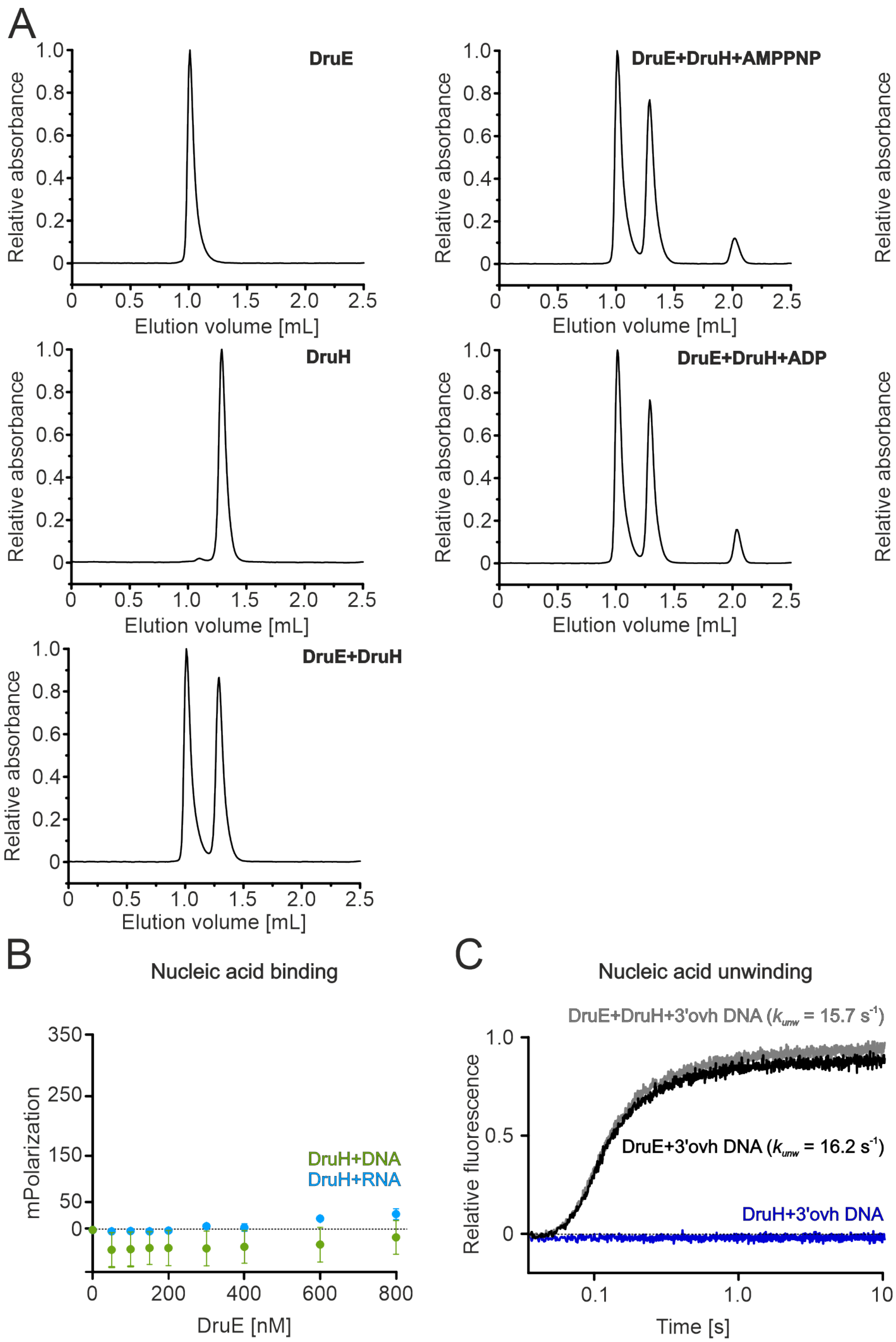

**Figure S11. DruE-DruH *in vitro* interaction studies. Related to Figure 6.**

(A) $UV_{280}$ traces of analytical SEC runs of the indicated samples.

(B) FA assays monitoring binding of DruH 15-nt ssDNA or ssRNA. Data represent the mean ± SEM of three independent experiments using the same biochemical samples.

(C) Unwinding of 3′ overhang (ovh) DNA by $DruE^{WT}$, DruH and $DruE^{WT}$ in the presence of DruH monitored *via* stopped-flow/fluorescence assays. Single representative traces of three independent measurements are shown. The unwinding rate of DruE differs from the values presented in figure 2-5 and S3 due to differences in buffer composition.

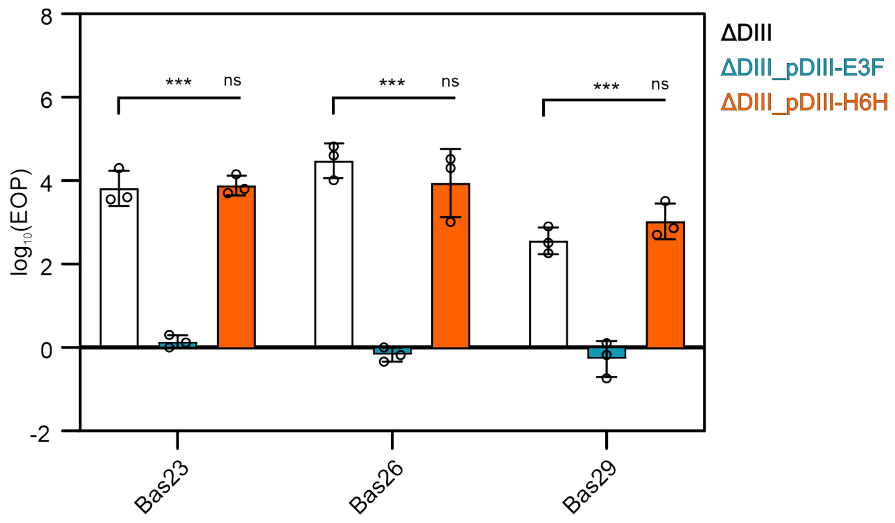

**Figure S12. Effect of affinity-tagged DruH and DruE on DIII mediated defense. Related to Figure 6.**

Efficiency of Plating (EOP) of Bas23, Bas26, Bas29 against *E. coli* ATCC8739 deleted of DIII (ΔDIII); *E. coli* ATCC8739 deleted of *druHE* and complemented with pDruHE_3F (p15a_DIII_E3F), coding for the DIII system under the native promoter, with a 3xFLAG tag at the C-terminus of DruE (ΔDIII_PDIII-E3F); or *E. coli* ATCC8739 deleted of DIII and complemented with p15a_DIII_H6H, coding for the DIII system under the native promoter, with a $His_6$-tag at the C-terminus of DruH (ΔDIII_PDIII-H6H). Bars and error bars represent the mean ± SD of three independent replicates. Open circles represent individual experiments. Significance was assessed by Benjamini, Krieger, and Yekutieli multiple unpaired t-tests. *, q ≤ 0.05; **, q ≤ 0.01; ***, q ≤ 0.001; ns, not significant.

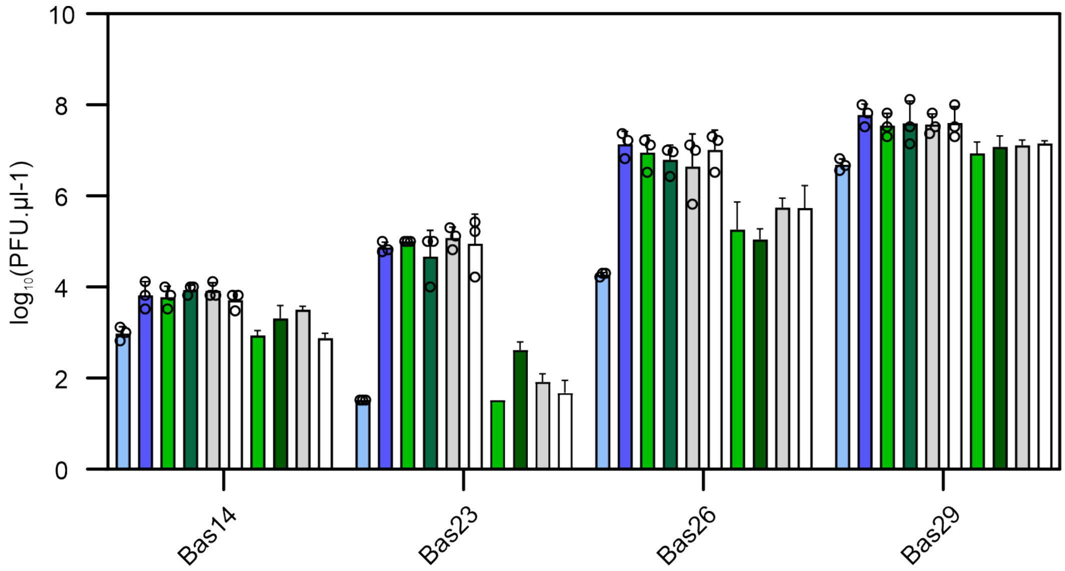

**Figure S13. Effects of *serA*, *rcdB*, *hns*, and *stpA* on DIII-mediated defense. Related to Figure 6.**

Phage titers of Bas14, Bas23, Bas26, and Bas29 against *E. coli* ATCC8739 WT (light blue), the single deletion mutant *E. coli* ATCC8739 Δ*druHE* (blue), the single deletion mutants *E. coli* ATCC8739 *ΔserA/rcdB/hns/stpA*, and the double deletions mutants *E. coli* ATCC8739 *ΔdruHE ΔserA/rcdB/hns/stpA* (respectively light green, green, grey and white). Bars and error bars represent the mean ± SD of three independent replicates. Open circles represent individual replicates.

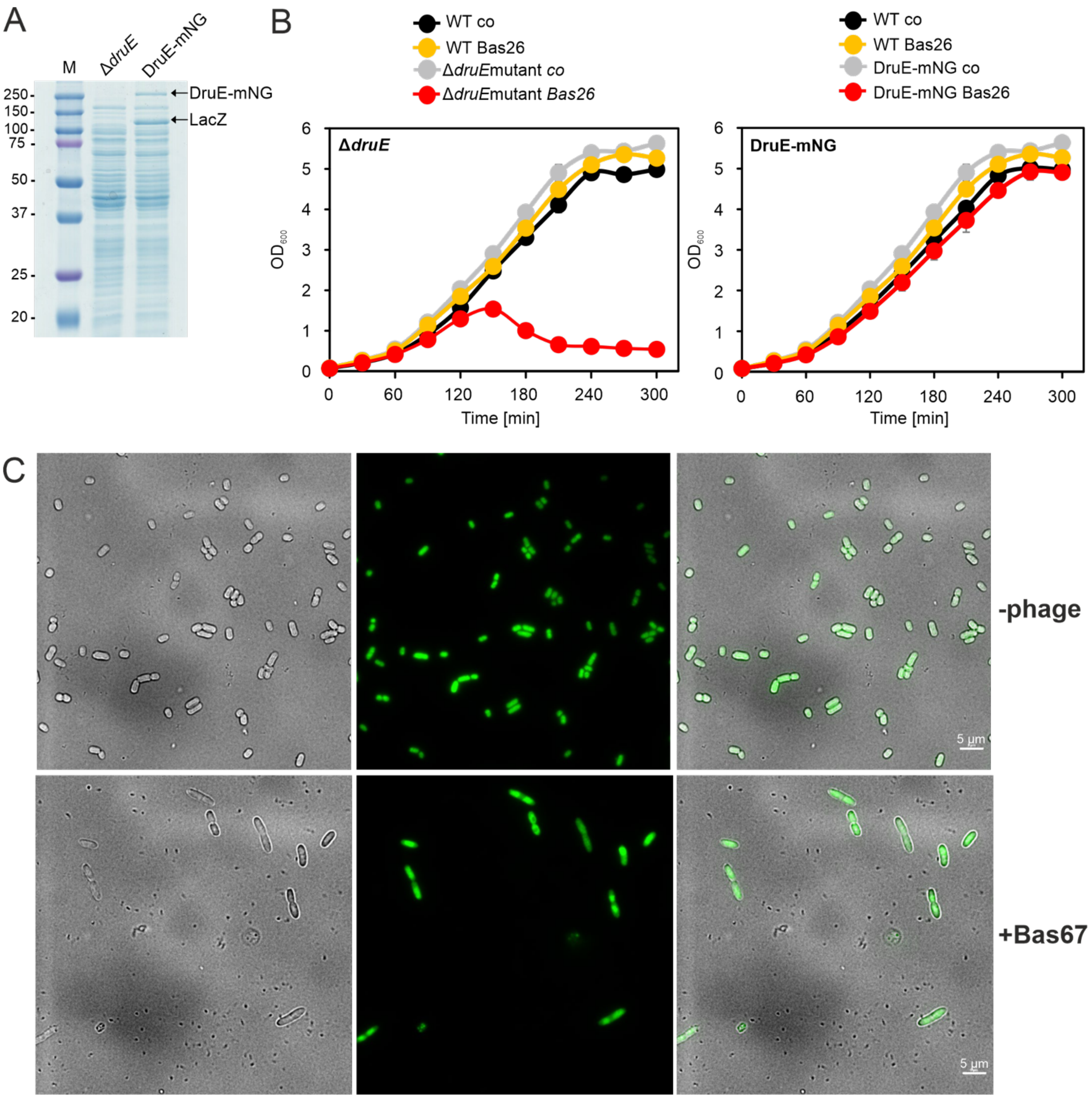

**Figure S14. Production of DruE-mNeonGreen and activity in phage defense. Related to Figure 3.**

(A) The EHEC strains EDL933 Δ*druE* and Δ*druE* pWKS30-*druE*-mNeonGreen were cultivated in LB with 0.2 mM IPTG and harvested after overnight growth followed by extraction of the intracellular proteins. The protein extract was separated using 12 % SDS-PAGE, the DruE-mNeonGreen and LacZ bands were cut and in-gel digested by trypsin. Tryptic peptides were subjected to LC-MS/MS analysis. Mascot search results identified the full-length DruE-mNeonGreen fusion protein and also the LacZ protein of EHEC strain, induced by IPTG. The full Mascot search results of DruE-mNeonGreen and LacZ are available at https://box.fu-berlin.de/s/4Sp5GKFBscALqwx.

(B) The EHEC strain encoding the DruE-mNeonGreen fusion restores the growth of the Δ*druE* mutant after infection by Bas26 at MOI 1, indicating functionality of the DruE-mNeonGreen fusion in phage defense in the EHEC strain. The EHEC strains EDL933 WT, Δ*druE*, and Δ*druE* pWKS30-*druE*-mNeonGreen were grown in LB to an $OD_{600}$ of 0.08, subjected to infection by phage Bas26 at MOI 1, and bacterial growth was monitored. Data represent means ± SD of three biological replicates.

(C) The EHEC strain EDL933 Δ*druE* pWKS30-*druE*-mNeonGreen was inoculated at $OD_{600}$ of 0.3 in LB with 0.2 mM IPTG and harvested after 2 h of growth without phage infection, and 2 h after infection by phage Bas67 at MOI 100. The EHEC cells were prepared and analyzed by TIRF microscopy as described in the Methods. Panels show transmitted light images (left), fluorescent images (center), and overlays (right).

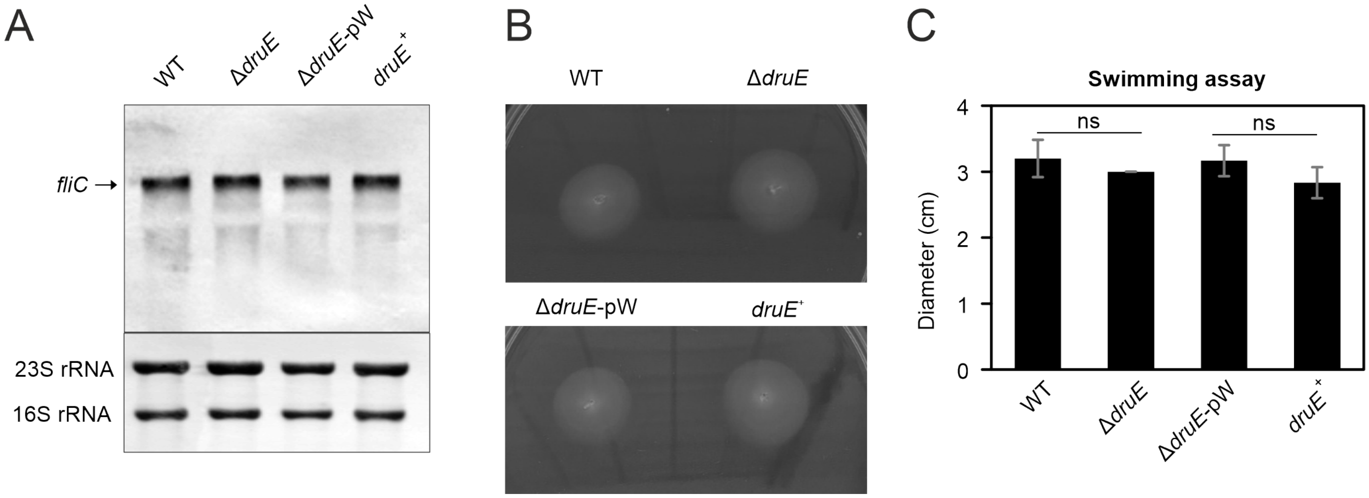

**Figure S15**. **Effect of DruE on *fliC* expression and EHEC motility. Related to Figure 1.**

(A) Northern blot analysis of *fliC* transcription. RNA was isolated from the indicated EHEC strains cultivated in LB medium and harvested at $OD_{600}$ = 0.8. Transcription of *fliC* was analyzed using Northern blots with a *fliC*-specific antisense RNA probe. Bands representing 16S and 23S rRNAs were stained with methylene blue as loading controls. Representative gels of three biological replicates are shown.

(B) Swimming assay testing the motility of the indicated EHEC strains. WT, EDL933 Δ*stx1/2*; Δ*druE*, EDL933 Δ*stx1/2* Δ*druE* mutant; *druE*+, *druE*-complemented EDL933 Δ*stx1/2* Δ*druE* mutant; Δ*druE*-pW, EDL933 Δ*stx1/2* Δ*druE* mutant transformed with empty pWKS30 plasmid. Overnight cultures (3 µl at $OD_{600}$ = 1) were spotted on LB plates containing 0.3 % soft agar.

(C) Swimming zones were quantified after incubation at 37 °C for 12 h. Values represent means of diameters of the swimming zones ± SD of three biological replicates.

**Table S1. CryoEM data collection and refinement statistics.**

| | **DruE monomer** (PDB 9TU7) (EMD-56251) | **DruE, dimer 1** (PDB 9TU8) (EMD-56252) | **DruE, dimer 2** (PDB 9TU9) (EMD-56253) | **DruE, dimer 3** (PDB 9TUA) (EMD-56254) | **DruE, dimer 4** (PDB 9TUB) (EMD-56255) | **DruE, dimer 5** (PDB 9TUC) (EMD-56256) | **DruH** (PDB 9TUD) (EMD-56257) |
|---|---|---|---|---|---|---|---|
| **Data collection and processing** | | | | | | | |
| Microscope | FEI Titan Krios G3i | | | | | | |
| Voltage [keV] | 300 | | | | | | |
| Camera | Falcon 3EC | | | | | | |
| Magnification (nominal/calibrated) | 96,000 | | | | | | 120,000 |
| Pixel size at detector [Å/pixel] | 0.832 | | | | | | 0.657 |
| Total electron exposure [$e^-/Å^2$] | 40 | | | | | | 42 |
| Exposure rate [$e^-$/pixel/s] | 0.7 | | | | | | 0.6 |
| No. of frames / exposure | 33 | | | | | | 32 |
| Defocus range [μm] | 0.8 – 2 | | | | | | |
| Automation software | EPU | | | | | | |
| Micrographs collected | 2,724 | | | | | | 1,950 |
| Micrographs used | 2,723 | | | | | | 1,863 |
| Total extracted particles | 660,861 | | | | | | 1,193,665 |
| Refined particles | 393,252 | | | | | | 67,814 |
| Final particles | 232,203 | 20,914 | 27,507 | 25,112 | 21,652 | 21,364 | 67,814 |
| Point-group or helical symmetry parameters | C1 | | | | | | |
| Global resolution [Å] | 2.9 | 3.9 | 3.6 | 3.7 | 3.8 | 4.1 | 3.3 |
| $FSC_{0.143}$ (unmasked/masked)[a] | 3.9 / 2.9 | 7.4 / 3.9 | 6.6 / 3.6 | 6.8 / 3.7 | 7.1 / 3.8 | 7.4 / 4.1 | 3.8 / 3.3 |
| Local resolution range [Å] | 2.5 - 26.5 | 3.2 - 52.5 | 3.1 - 53 | 3.1 - 52.6 | 3.2 - 52.4 | 3.4 - 56.7 | 2.5 - 35 |
| Map sharpening *B* factor [$Å^2$] / (*B* factor range) | -98 | -59.4 | -70.8 | -71.4 | -67.4 | -69.1 | -84 |
| Map sharpening methods | Local resolution | | | | | | |

| Refinement package | PHENIX (1.21.2-5419) real.space.refine | | | | | | |
|---|---|---|---|---|---|---|---|
| **Model composition** | | | | | | | |
| Non-hydrogen atoms | 16855 | 34015 | 33663 | 31858 | 31776 | 32402 | 8.946 |
| Protein residues (subunit 1 / 2) | 2003 | 2068 / 2050 | 2075 / 2050 | 2046 / 1823 | 1843 / 2048 | 2049 /1833 | - |
| DNA nucleotides (subunit 1 / 2) | 43 | 53 / 7 | 41 / 0 | 12 / 41 | 35 / 5 | 36 / 39 | - |
| ATPγS (subunit 1 / 2) | 1 / - | 1 / 1 | 1 / 1 | 1 / 1 | 1 / 1 | 1 / - | - |
| $Zn^{2+}$ ions (subunit 1 / 2) | 5 / - | 5 / 5 | 5 / 5 | 5 / 5 | 5 / 5 | 5 / 5 | - |
| **Model Refinement** | | | | | | | |
| **Model-Map scores** | | | | | | | |
| CC (mask)[b] | 0.87 | 0.82 | 0.77 | 0.75 | 0.83 | 0.78 | 0.77 |
| CC (volume) | 0.87 | 0.82 | 0.77 | 0.74 | 0.83 | 0.77 | 0.77 |
| Average FSC (masked/unmasked) | 2.5/2.5 | 3.8/3.9 | 3.4/3.6 | 3.6/3.7 | 3.5/3.8 | 4.1/4.1 | 2.7/2.8 |
| **Average B factors [Å²]** | | | | | | | |
| Overall | 128 | 261 | 217 | 219 | 207 | 252 | 83 |
| Protein residues (subunit 1 / 2) | 127 / - | 242 / 282 | 165 / 266 | 199 / 247 | 221 / 186 | 221 / 273 | |
| DNA nucleotides (subunit 1 / 2) | 140 / - | 251 / 166 | 291 / - | 100 / 157 | 408 / 127 | 394 / 400 | - |
| ATPγS (subunit 1 / 2) | 140 / - | 231 / 275 | 152 / 268 | 238 / 279 | 244 / 190 | 233 / - | - |
| $Zn^{2+}$ ions (subunit 1 / 2) | 145 / - | 233 / 337 | 165 / 398 | 284 / 274 | 250 / 223 | 282 / 287 | - |
| **RMSD[c] from ideal values** | | | | | | | |
| Bond lengths [Å] | 0.004 | 0.002 | 0.006 | 0.003 | 0.003 | 0.003 | 0.003 |
| Bond angles [°] | 0.557 | 0.565 | 0.725 | 0.658 | 0.597 | 0.606 | 0.656 |
| **Validation** | | | | | | | |
| MolProbity score | 1.71 | 2.18 | 2.77 | 2.30 | 2.09 | 2.23 | 2.48 |
| CaBLAM outliers [%] | 2.0 | 2.8 | 3.1 | 2.8 | 2.9 | 2.8 | 4.07 |
| Clashscore | 8.5 | 14.9 | 25.9 | 16.1 | 14.6 | 18.4 | 13.05 |
| Poor rotamers [%] | 1.2 | 1.7 | 4.0 | 1.9 | 1.4 | 1.7 | 4.55 |

| | | | | | | | |
|---|---|---|---|---|---|---|---|
| Cβ deviations | 0 | 0 | 0 | 0 | 0 | 0 | 0 |
| **Ramachandran plot** | | | | | | | |
| Favored [%] | 96.8 | 95.4 | 93.8 | 95.4 | 95.6 | 95.6 | 94.9 |
| Allowed [%] | 3.2 | 4.6 | 6.2 | 4.5 | 4.4 | 4.3 | 5 |
| Outliers [%] | 0.1 | 0.0 | 0.0 | 0.1 | 0.0 | 0.0 | 0.1 |
| **Ramachandran plot *Z*-score (RMSD)** | | | | | | | |
| Whole | 0.20 (0.19) | -0.39 (0.13) | -1.14 (0.13) | -0.40 (0.14) | -0.18 (0.14) | -0.20 (0.14) | -0.34 (0.26) |
| Helix | 1.53 (0.20) | 1.35 (0.14) | 0.33 (0.13) | 0.99 (0.14) | 1.29 (0.14) | 1.14 (0.14) | 0.87 (0.29) |
| Sheet | -0.27 (0.28) | -0.53 (0.21) | -1.12(0.21) | -0.43 (0.22) | -0.30 (0.21) | -0.57 (0.22) | -0.14 (0.38) |
| Loop | -0.81 (0.19) | -1.49 (0.14) | -1.55 (0.14) | -1.27 (0.14) | -1.28 (0.14) | -1.03 (0.14) | -0.92 (0.26) |

[a] FSC, Fourier shell correlation
[b] CC, correlation coefficient
[c] RMSD, root mean square deviation

**Table S2. Mass spectrometry analyses.** See the attached .csv file.

**Table S3. Bacterial strains and plasmids used in this study.**

| *Escherichia coli* strain | Description | Source |
|---|---|---|
| DH5α | F-φ80dlacZ Δ(lacZYA-argF) U169 deoRsupE44ΔlacU169 (f80lacZDM15) hsdR17 recA1 endA1 (rk- mk+) supE44gyrA96 thi-1 gyrA69 relA1 | [1] |
| High Five cells | *Trichoplusia ni* cell line commonly used for protein expression | Thermo Scientific |
| Sf9 insect cells | | Gibco |
| BL21 STAR (DE3) | | Invitrogen |
| K-12 MG1655 ΔRM | K-12 MG1655 ΔRM | [2] |
| K-12 MG1655 | N/A | Strain from Susan Gottesman's lab |
| K-12 MG1655 p15a_empty | K-12 MG1655 with only the vector backbone | This study |
| K-12 MG1655 p15a_DTIII-A | K-12 MG1655 expressing Druantia type III-A operon under its putative native promoter on a low copy plasmid | This study |
| K-12 MG1655 p15a_DTIII-A_Δ*druE* | K-12 MG1655 expressing Druantia type III-A operon deleted of the *druE* gene | This study |
| K-12 MG1655 p15a_DTIII-A_Δ*druH* | K-12 MG1655 expressing Druantia type III-A operon deleted of the *druH* gene | This study |
| K-12 MG1655 p15a_DTIII-A_Δ*druH* + p15a_DTIII-A_Δ*druE* | K-12 MG1655 expressing Druantia type III-A operon deleted of the *druE* gene complemented with a p15a_DTIII-A_Δ*druH* | This study |
| K-12 MG1655 p15a_DTIII-A_G118V | K-12 MG1655 expressing Druantia type III-A operon with DruE E252A substitution | This study |
| K-12 MG1655 p15a_DTIII-A_G120V | K-12 MG1655 expressing Druantia type III-A operon with DruE G120V substitution | This study |
| K-12 MG1655 p15a_DTIII-A_D251A | K-12 MG1655 expressing Druantia type III-A operon with DruE D251A substitution | This study |
| K-12 MG1655 p15a_DTIII-A_E252A | K-12 MG1655 expressing Druantia type III-A operon with DruE E252A substitution | This study |
| K-12 MG1655 p15a_DTIII-A_R1135L | K-12 MG1655 expressing Druantia type III-A operon with DruE R1135L substitution | This study |

| | | |
|---|---|---|
| K-12 MG1655 p15a_DTIII-A_R1138L | K-12 MG1655 expressing Druantia type III-A operon with DruE R1138L substitution | This study |
| K-12 MG1655 p15a_DTIII-A_C1681G | K-12 MG1655 expressing Druantia type III-A operon with DruE C1681G substitution | This study |
| K-12 MG1655 p15a_DTIII-A_C1683G | K-12 MG1655 expressing Druantia type III-A operon with DruE C1683G substitution | This study |
| K-12 MG1655 p15a_DTIII-A_C1687G | K-12 MG1655 expressing Druantia type III-A operon with DruE C1687G substitution | This study |
| EDL933 Δ*stx1/2* | EDL933 *stx1/2* deletion mutant | [3] |
| EDL933 Δ*stx1/2* Δ*druE* | EDL933 *stx1/2 druE* (z5898) scar mutant | [4] |
| EDL933 Δ*stx1/2* Δ*druE*-pWKS30 | EDL933 *stx1/2 druE* (z5898) scar mutant complemented with pWKS30 (empty plasmid) | [4] |
| EDL933 Δ*stx1/2* Δ*druE*-pWKS30-*druE* | EDL933 *stx1/2 druE* (z5898) scar mutant complemented with pWKS30-*druE* | [4] |
| EDL933 Δ*stx1/2* Δ*druE*-pWKS30-*druE*-(1-1171) | EDL933 *stx1/2 druE* (z5898) scar mutant complemented with pWKS30-*druE* (1-1171) | This study |
| EDL933 Δ*stx1/2* Δ*druE*-pWKS30-*druE*-(1-1502) | EDL933 *stx1/2 druE* (z5898) scar mutant complemented with pWKS30-*druE* (1-1502) | This study |
| EDL933 Δ*stx1/2* Δ*druE*-pWKS30-*druE*-(1-1723) | EDL933 *stx1/2 druE* (z5898) scar mutant complemented with pWKS30-*druE* (1-1723) | This study |
| EDL933 Δ*stx1/2* Δ*druE*-pWKS30-*druE*-(1570-2104) | EDL933 *stx1/2 druE* (z5898) scar mutant complemented with pWKS30-*druE*-Cterm | This study |
| EDL933 Δ*stx1/2* Δ*druE*-pWKS30-*druE*-ΔDD | EDL933 *stx1/2 druE* (z5898) scar mutant complemented with pWKS30-*druE* | This study |
| EDL933 Δ*stx1/2* Δ*druE*-pWKS30-*druE*-Δlock | EDL933 *stx1/2 druE* (z5898) scar mutant complemented with pWKS30-*druE*-Δlock | This study |
| EDL933 Δ*stx1/2* Δ*druE*-pWKS30-*druE* W1480A | EDL933 *stx1/2 druE* (z5898) scar mutant complemented with pWKS30-*druE*-W1480A | This study |
| EDL933 Δ*stx1/2* Δ*druE*-pWKS30-*druE*-F1327A | EDL933 *stx1/2 druE* (z5898) scar mutant complemented with pWKS30-*druE*-F1327A | This study |
| EDL933 Δ*stx1/2* Δ*druE*-pWKS30-*druE*-CC1687,1690AA | EDL933 *stx1/2 druE* (z5898) scar mutant complemented with pWKS30-*druE*-C1687,1690A | This study |
| EDL933 Δ*stx1/2* Δ*druE*-pWKS30-*druE*-CC1681,1683AA | EDL933 *stx1/2 druE* (z5898) scar mutant complemented with pWKS30-*druE*-C1681,1683A | This study |
| EDL933 Δ*stx1/2* Δ*druE*-pWKS30-*druE*-E252A | EDL933 *stx1/2 druE* (z5898) scar mutant complemented with pWKS30-*druE*-E252A | This study |
| EDL933 Δ*stx1/2* Δ*druE*-pWKS30-*druE*-mNeonGreen | EDL933 *stx1/2 druE* (z5898) scar mutant complemented with pWKS30-*druE*-mNeonGreen | This study |

| | | |
|---|---|---|
| ATCC8739 | Wildtype strain | [5] |
| ATCC8739 Δ*druHE* | ATCC8739 *druHE* deletion mutant | [5] |
| ATCC8739 Δ*rcdB* | ATCC8739 *rcdB* deletion mutant | This study |
| ATCC8739 Δ*serA* | ATCC8739 *serA* deletion mutant | This study |
| ATCC8739 Δ*stpA* | ATCC8739 *stpA* deletion mutant | This study |
| ATCC8739 Δ*hns* | ATCC8739 *hns* deletion mutant | This study |
| ATCC8739 Δ*druHE* Δ*rcdB* | ATCC8739 *druHE rcdB* deletion mutant | This study |
| ATCC8739 Δ*druHE* Δ*serA* | ATCC8739 *druHE serA* deletion mutant | This study |
| ATCC8739 Δ*druHE* Δ*stpA* | ATCC8739 *druHE stpA* deletion mutant | This study |
| ATCC8739 Δ*druHE* Δ*hns* | ATCC8739 *druHE hns* deletion mutant | This study |
| ATCC8739 Δ*druHE* p15a_DTIII-A_E3F | ATCC8739 *druHE* deletion mutant complemented with p15a_DTIII-A_E3F | This study |
| ATCC8739 Δ*druHE* p15a_DTIII-A_3FE | ATCC8739 *druHE* deletion mutant complemented with p15a_DTIII-A_3FE | This study |
| ATCC8739 Δ*druHE* p15a_DTIII-A_H6F | ATCC8739 *druHE* deletion mutant complemented with p15a_DTIII-A_H6F | This study |
| ATCC8739 Δ*druHE* p15a_DTIII-A_6FH | ATCC8739 *druHE* deletion mutant complemented with p15a_DTIII-A_6FH | This study |
| ATCC8739 Δ*druHE* p15a_DTIII-A | ATCC8739 Δ*druHE* p15a_DTIII-A complemented with p15a_DTIII-A | [5] |
| **Plasmid** | **Description** | **Reference** |
| pWKS30 | $Amp^R$, *lacZ*α, pSC101 *ori*, f1 *ori*, MCS | [6] |
| pWKS30-*druE* | pWKS30 expressing *druE* under *lac*-promoter | [4] |
| pWKS30-*druE*-(1-1171) | pWKS30 expressing *druE*-1-1171 under *lac*-promoter | This study |
| pWKS30-*druE*-(1-1502) | pWKS30 expressing *druE*-1-1502 under *lac*-promoter | This study |
| pWKS30-*druE*-(1-1723) | pWKS30 expressing *druE*-1-1723 under *lac*-promoter | This study |
| pWKS30-*druE*-(1570-2104) | pWKS30 expressing *druE*-1520-2104 under *lac*-promoter | This study |
| pWKS30-*druE*-ΔDD | pWKS30 expressing *druE*-ΔDD under *lac*-promoter | This study |
| pWKS30-*druE*-Δlock | pWKS30 expressing *druE*-Δlock under *lac*-promoter | This study |
| pWKS30-*druE*-W1480A | pWKS30 expressing *druE*-W1480A under *lac*-promoter | This study |

| | | |
|---|---|---|
| pWKS30-*druE*-F1327A | pWKS30 expressing *druE*-F1327A under *lac*-promoter | This study |
| pWKS30-*druE*-CC1687/1690AA | pWKS30 expressing *druE*-CC1687,1690A under *lac*-promoter | This study |
| pWKS30-*druE*-CC1681/1683AA | pWKS30 expressing *druE*-CC1681,1683A under *lac*-promoter | This study |
| pWKS30-*druE*-E252A | pWKS30 expressing *druE*-E252A under *lac*-promoter | This study |
| pFL | $Amp^R$, $Gen^R$, f1 *ori*, MCS, polyhedron and p10 promotors, 10His, TEV site | [7] |
| pFL-*druE*-(1-1171) | pFL expressing *druE*-1-1171 under the polyhedron promotor | This study |
| pFL-*druE*-(1-1502) | pFL expressing *druE*-1-1502 under the polyhedron promotor | This study |
| pFL-*druE*-(1-1723) | pFL expressing *druE*-1-1723 under the polyhedron promotor | This study |
| pFL-*druE*-(1570-2104) | pFL expressing *druE*-1570-2104 under the polyhedron promotor | This study |
| pFL-*druE*-ΔDD | pFL expressing *druE*-ΔDD under the polyhedron promotor | This study |
| pFL-*druE*-Δlock | pFL expressing *druE*-Δlock under the polyhedron promotor | This study |
| pFL-*druE*-W1480A | pFL expressing *druE*-W1480A under the polyhedron promotor | This study |
| pFL-*druE*-F1327A | pFL expressing *druE*-F1327A under the polyhedron promotor | This study |
| pFL-*druE*-CC1687,1690AA | pFL expressing *druE*-CC1687,1690A under the polyhedron promotor | This study |
| pFL-*druE*-CC1681,1683AA | pFL expressing *druE*-CC1681,1683A under the polyhedron promotor | This study |
| pFL-*druE*-E252A | pFL expressing *druE*-E252A under the polyhedron promotor | This study |
| pWKS30-*druE*-mNeonGreen | pWKS30 expressing *druE*-mNeonGreen under *lac*-promoter | This study |
| pEM8731 | pEM8731 (pKH70-$P_{rpsM}$-*mNeonGreen*, AmpR) | [8] |
| pETM11 | $Kan^R$, f1 *ori*, MCS, T7 and lacI promotor, 6His, TEV site | EMBL Heidelberg |
| pETM11-*druH* | pETM11 expressing *druH* under the T7 promotor | This study |
| p15a_empty | $Cm^R$, p15a *ori*, ampicillin promoter. Empty-vector control. | This study |

| | | |
|---|---|---|
| pDruHE | p15a_empty expressing Druantia type III-A from ATCC8739 under its native promoter | [5] |
| pDruH | pDruHE with *druE* deleted | This study |
| pDruE | pDruHE with *druH* deleted; $Amp^R$ | This study |
| pDruHE_G118V | pDruHE with DruE G118V substitution | This study |
| pDruHE_G120V | pDruHE with DruE G120V substitution | This study |
| pDruHE_D251A | pDruHE with DruE D251A substitution | This study |
| pDruHE_E252A | pDruHE with DruE E252A substitution | This study |
| pDruHE_R1135L | pDruHE with DruE R1135L substitution | This study |
| pDruHE_R1138L | pDruHE with DruE R1138L substitution | This study |
| pDruHE_C1681G | pDruHE with DruE C1681G substitution | This study |
| pDruHE_C1683G | pDruHE with DruE C1683G substitution | This study |
| pDruHE_C1687G | pDruHE with DruE C1687G substitution | This study |
| pDruHE_E3F | pDruHE with a 3X-FLAG tag on the C-terminus of DruE | This study |
| pDruHE_3FE | pDruHE with a 3X-FLAG tag on the N-terminus of DruE | This study |
| pDruHE_H6F | pDruHE with a 6 Histidines tag on the N-terminus of DruH | This study |
| pDruHE_6FH | pDruHE with a 6 Histidines tag on the C-terminus of DruH | This study |

$Amp^R$, ampicillin resistance; $Gen^R$, gentamicin resistance; $Kan^R$, kanamycin resistance; $Cm^R$; chloramphenicol resistance

**Table S4. Oligonucleotides used in this study.** FP, forward primer; RP, reverse primer; SDM, site-directed mutagenesis.

| Name | Sequence (5′-to-3′) | Source |
|---|---|---|
| **Primers for cloning gene block 1 into pFL vector** | | |
| pFL_EcoRI_FP | GGTTCAGCAGAGGGGCGTAGTGCAGCTGCTGAACGTC CATGAATTCGGCGCCCTGAAAATAAAGATTC | Eurofins Genomics |
| pFL_block1_RP | CTATCGACGCCCTGAAGAAAGCTATGGCCGCCAACGAA GAACAAGCTTGTCGAGAAGTACTAGAGGATC | |
| block1_FP | TTCCATGGACGTTCAGCAGCTGCAC | |
| block1_RP | GTTCTTCGTTGGCGGCCATAGCTTTC | |
| **Primers for cloning gene block 2 and 3 into block 1-containing pFL vector** | | |
| block2_FP | GTCCTGCCTCTGAAGAACTCTATCGACGC | Eurofins Genomics |
| block3_RP | CTTATCCGGCAATGATGTAAGACCT | |
| pFL-block1_RP | GTTCTTCGTTGGCGGCCATAGCTTTC | |
| pFL-block1_FP | CTCCTGGAAACGACCTGAGGTCTTACATCATTGCCGGA TAAAAGCTTGTCGAGAAGTACTAGAGGATC | |
| **Primers for cloning pETM11-*druH*** | | |
| pETM11_*druH*_NcoI | AGCCATGGAAACTCCAGTGAACCCGCTG | Eurofins Genomics |
| pETM11_*druH*_HindII | AGAAGCTTTTAGTGGCACACGCCCGC | |
| **Primers for introducing point mutations and truncations *via* inverse PCR** | | |
| *druE*-(1-1171)_FP | TAAAAGCTTGTCGAGAAGTACTAGAGGATC | Eurofins Genomics |
| *druE*-(1-1171)_RP | GGCAGTGGTGTTGAAAGCCCAC | |
| *druE-(*1-1502)_FP | TAAAAGCTTGTCGAGAAGTACTAGAGGATC | |
| *druE-(*1-1502)_RP | GCGTCCCAGTCCGGCATCT | |
| *druE*-(1570-2104)_FP | TGCCACGGACACGTCCACAAG | |
| *druE*-(1570-2104)_RP | GAATTCGGCGCCCTGAAAATAAAGAT | |
| *druE*-Δlock_(1-509_GSG_527-2104)_FP | CTGGCGACGAACTGGACAACAACTCCCA | |

| | | |
|---|---|---|
| *druE*_Δlock(1-509_GSG_527-2104)_RP | AGCCAGCAGCGGCAGGTTGAG | |
| *druE*-ΔDD(1-676_GSG_760-2104)_ FP | TGGCAAGACCATGTCCTGGCAGACCCTG | |
| *druE*-ΔDD(1-676_GSG_760-2104)_RP | GAGCCGTTGGCCAGCTCTTGGTAGCACAG | |
| *druE*-F1327A_FP | GCCCCTGTCGGAGTCGTGAAC | |
| *druE*-F1327A_RP | GCAGGCAGCTGAGCAGG | |
| *druE*-W1480A_FP | GAAGGACCCAAGAGTCTTGGTC | |
| *druE*-W1480A_RP | GCAGGCAGCTGAGCAGG | |
| *druE*-(1-1723)_FP | TTAAGCGAGGTTCAGGCGCTGC | |
| *druE*-(1-1723)_RP | AAGCTTGTCGAGAAGTACTAGAGGATCA | |
| *druE*$^{CC1681/1683AA}$_FP | CAGAAAGCTGCTCGACGCCTCTGCCGACAAGTTCTGTCAC | |
| *druE*$^{CC1681/1683AA}$_RP | GTGACAGAACTTGTCGGCAGAGGCGTCGAGCAGCTTTCTG | |
| **Vectors synthesized/cloned by GenScript** | | |
| pFL-*druE*$^{E252A}$ | | |
| pFL-*druE*$^{CC1687/1690AA}$ | | |
| **Primers for amplifying *the* pWKS30 backbone for *druE* variants** | | |
| pWKS30_FP | AGCTGTTTCCTGTGTGAAATTGTT | Eurofins Genomics |
| pWKS30_RP | ACCATGATTACGCCAAGCGC | |
| **Primers for amplifying the pWKS30 backbone for pWKS30-*druH*** | | |
| pWKS30 for *druH*_FP | CAAGCAGGAGTATGTCATTGAATGATTACGCCAAGCGC | Eurofins Genomics |
| pWKS30 for *druH*_RP | GTGGGTTAACTGGAGTTTTCACAGCTGTTTCCTGTGTGAAATTG | |
| **Primers for amplifying *druH* from the EDL933 genome** | | |
| *druH*_FP | GTGAAAACTCCAGTTAACCCAC | Eurofins Genomics |
| *druH*_RP | TCAATGACATACTCCTGCTTGT | |
| **Primers for amplifying *druE* variants from the pFL vectors for insertion into pWKS30** | | |

| | | |
|---|---|---|
| pW_*druE*_FP | CAATTTCACACAGGAAACAGCTATGGACGTTCAGCAGC TGC | Eurofins Genomics |
| pW_*druE*_RP | CAATTTCACACAGGAAACAGCTATGGACGTTCAGCAGC TGC | |
| pW_*druE*-(1-1171)_RP | GCT TGG CGT AAT CAT GGT TTA GGC AGT GGT GTT GAA AGC C | |
| pW_*druE*-(1-1502)_RP | CGC TTG GCG TAA TCA TGG TTT AGC GTC CCA GTC CGG CAT C | |
| pW_*druE*-(1570-2104)_FP | CAA TTT CAC ACA GGA AAC AGC TAT GTG CCA CGG ACA CGT CC | |
| pW_*druE*-(1-1723)_RP | ATTGCGCGCTTGGCGTAATCATGGTTTAAGCGAGGTTC AGGCGCTG | |
| pWKS-*druE*-FP | CAATTTCACACAGGAAACAGCTATGGACGTTCAGCAACT GCATT | Eurofins Genomics |
| *druE*-neoGreen-RP | TCCTCGCCCTTCGATACCATGCTGGCGCTGCCGGCGCT TCCAGCGATAATATAACTCC | |
| neoGreen-FP | GGAGTTATATTATCGCTGGAAGCGCCGGCAGCGCCAGC ATGGTATCGAAGGGCGAGGA | |
| neoGreen-RP | GCGCGCTTGGCGTAATCATGGTTCATTTATACAGTTCAT CCATGCC | |
| **Oligonucleotides for RNA/DNA binding assays** | | |
| FAM-DNASS15 | [5-FAM]-GCTGCCAGACCAAAT | IBA |
| FAM-RNASS15 | [5-FAM]-GCUGCCAGACCAAAU | |
| **Oligonucleotide for ATPase assays** | | |
| RNA_U_15 | UUUUUUUUUUUUUUU | IBA |
| DNA_19nts | GCGTCCCAGTCCGGCATCT | |
| **Oligonucleotides for unwinding assays** | | |
| DNA_3'ovh | [Atto540Q]-GGCCGCGAGCCGGAAATTTAATTATAAACCAGACCGTC TCCTC<br>CGGCTCGCGGCC-[Alexa488] | IBA |
| DNA_5'ovh | CTCCTCTGCCAGACCAAATATTAATTTAAAGGCCGAGCG CCGG-[Atto540Q]<br>[Alexa488]-CCGGCGCTCGGC | |

| | | |
|---|---|---|
| RNA_3’ovh | [Atto540Q]-<br>GGCCGCGAGCCGGAAAUUUAAUUAUAAACCAGACCGU<br>CUCCUC<br>CGGCUCGCGGCC-[Alexa488] | |
| DNA_blocked_ovh | [Atto540Q]-<br>GGCCGCGAGCCGGAAATTTAATTATAAACCAGACCGTC<br>TCCTC-3’<br>CGGCTCGCGGCC-[Alexa488]<br>GAGGAGACGGTC | IBA |
| **Oligonucleotides for cryoEM** | | |
| Forked_DNA | ATCGATAGTCTCTAGGCTGCCAGACCAAAT<br>TAAACCAGACCGTCGGAAGAGACTATCGAT | Eurofins Genomics |
